# A projection specific logic to sampling visual inputs in mouse superior colliculus

**DOI:** 10.1101/272914

**Authors:** Katja Reinhard, Chen Li, Quan Do, Emily Burke, Steven Heynderickx, Karl Farrow

## Abstract

Using sensory information to trigger different behaviours relies on circuits that pass-through brain regions. However, the rules by which parallel inputs are routed to different downstream targets is poorly understood. The superior colliculus mediates a set of innate behaviours, receiving input from ~30 retinal ganglion cell types and projecting to behaviourally important targets including the pulvinar and parabigeminal nucleus. Combining transsynaptic circuit tracing with *in-vivo* and *ex-vivo* electrophysiological recordings we observed a projection specific logic where each collicular output pathway sampled a distinct set of retinal inputs. Neurons projecting to the pulvinar or parabigeminal nucleus uniquely sampled 4 and 7 cell types, respectively. Four others innervated both pathways. The visual response properties of retinal ganglion cells correlated well with those of their disynaptic targets. These findings suggest that projection specific sampling of retinal inputs forms a mechanistic basis for the selective triggering of visually guided behaviours by the superior colliculus.

## Introduction

The nervous system is built from a large set of diverse neuronal cell types that work together to process information and generate behavior (Zeng and Sanes, 2017). Sets of connected neurons can be divided up into “hard-wired” circuits that enable robust, stereotyped, reflex-like behavioral responses (Chen et al., 2011; Lorente de No, 1933; Lundberg, 1979), and flexible networks that modify their computations based on context and experience (Dhawale et al., 2017; Rose et al., 2016). Many innate behaviors rely on subcortical circuits involving the same sets of brain structures in different species (Aponte et al., 2011; Gandhi and Katnani, 2011; Hong et al., 2018; Tinbergen, 1951). In the visual system it remains unclear to what extent these circuits have hard-wired rules linking their inputs with downstream targets (Cruz-Martín et al., 2014; Ellis et al., 2016; Gale and Murphy, 2014, 2018; Glickfeld et al., 2013; Liang et al., 2018; Morgan et al., 2016; Rompani et al., 2017).

The output of the retina, the first stage of visual processing, consists of over 30 different ganglion cell types which can be distinguished by their dendritic anatomy, response properties, or molecular markers (Baden et al., 2016; Dhande et al., 2015; Farrow and Masland, 2011; Martersteck et al., 2017; Roska and Werblin, 2001; Sanes and Masland, 2014). Each ganglion cell type informs one or several brain areas about a certain feature of the visual world (Ellis et al., 2016; Martersteck et al., 2017). One of the major retinorecipient areas is the superior colliculus, which receives approximately 85% of the retinal outputs in rodents (Ellis et al., 2016; Hofbauer and Dräger, 1985; Linden and Perry, 1983; Vaney et al., 1981).

The rodent superior colliculus is a layered brain structure that receives inputs from all sensory modalities and targets various nuclei of the midbrain and brainstem. The superficial gray and the optic layer form the most dorsal layers of the superior colliculus and are primarily innervated by the retina (May, 2006). These visual layers consist of several groups of neurons with diverse morphology, visual response properties and long-range targets that include the lateral pulvinar, lateral geniculate nucleus and parabigeminal nucleus. Each neuron of the superficial superior colliculus has been estimated to receive input from on average six retinal ganglion cells (Chandrasekaran et al., 2007). However, the different ganglion cell types that provide input to specific superior collicular output pathways have not been characterized. As a result, it is unknown whether each output pathway of the superior colliculus shares a common or different set of retinal inputs, and consequently whether different visual inputs give rise to the different behaviors initiated by the colliculus (Evans et al., 2018; Shang et al., 2015, 2018; Wei et al., 2015; Zhang et al., 2019).

To determine the wiring rules underlying the integration of retinal information by different output pathways of the superior colliculus, we used a combination of transsynaptic viral tracing and molecular markers to specifically label the retinal ganglion cells at the beginning of two circuits: one targeting the parabigeminal nucleus (colliculo-parabigeminal circuit) and the second targeting the pulvinar (colliculo-pulvinar circuit). Using quantitative analysis of the retinal ganglion cell morphology and comparison of the visual response properties in the retina and target nuclei, we found strong specificity in the routing of visual information through the superior colliculus.

## Results

### Transsynaptic tracing of retinal ganglion cells from targets of the superior colliculus

To determine if visual features are selectively sampled by the two targeted output pathways of the mouse superior colliculus, we used rabies-based viral tools to label retinal ganglion cells innervating either the colliculo-parabigeminal or colliculo-pulvinar circuit. We injected the parabigeminal nucleus (Figure 1A-C) or lateral pulvinar (Figure S1A-C) with herpes-simplex virus (HSV) expressing rabies-G, TVA and mCherry, and subsequently injected EnvA-coated rabies virus coding for GCaMP6s (EnvA-SADΔG-GCaMP6s) into the superficial layers of the superior colliculus (see Methods). This transsynaptic viral infection strategy resulted in the expression of GCaMP6s in several dozen retinal ganglion cells per retina that specifically innervate the targeted circuit. As the lateral pulvinar lies adjacent to the lateral geniculate we used the Ntsr1-GN209-Cre mouse line to ensure specific infection of neurons projecting to the lateral pulvinar (Gale and Murphy, 2014).

**Figure 1.**
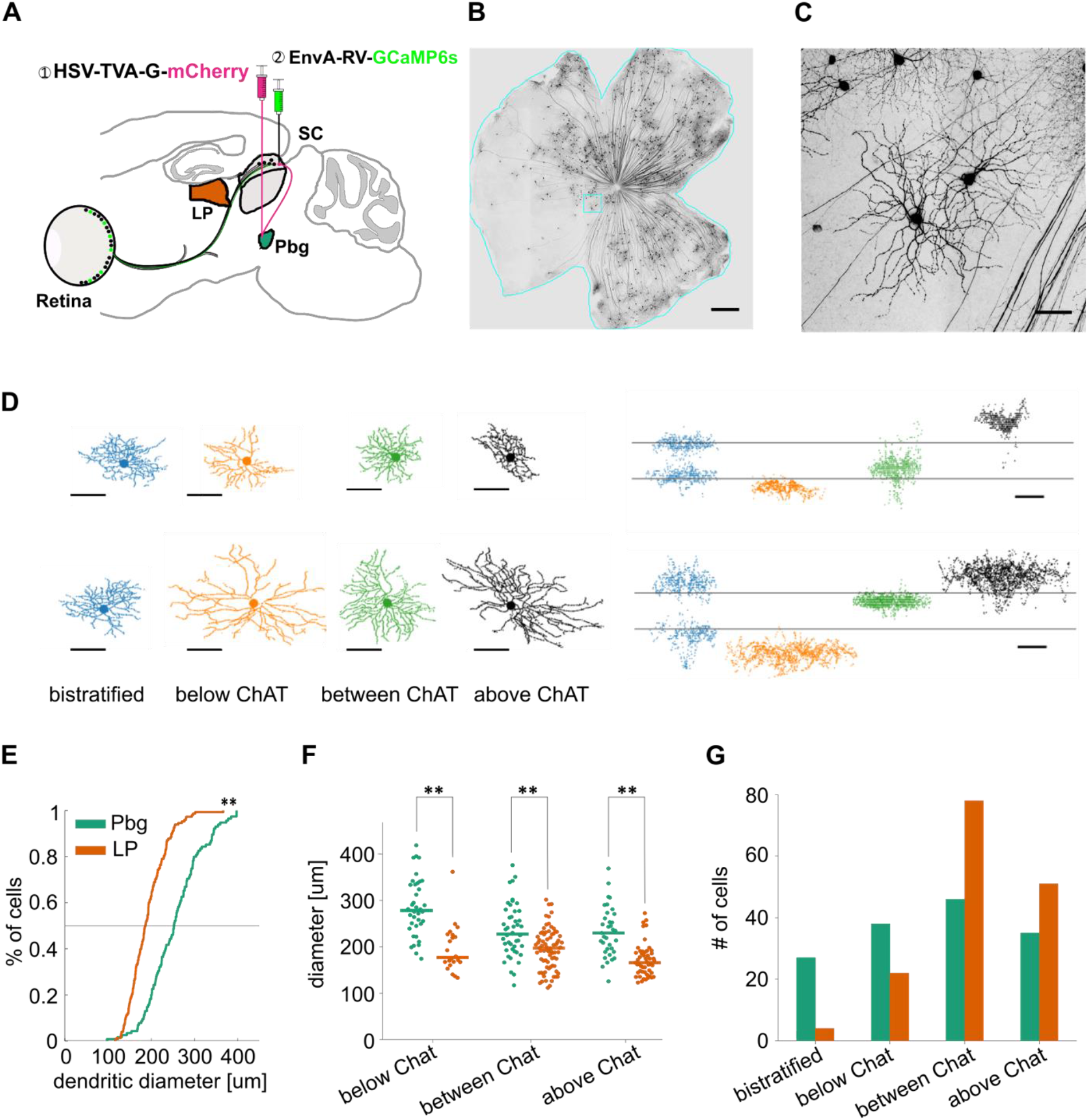
Transsynaptic tracing of retinal ganglion cells from the parabigeminal nucleus and the lateral pulvinar. **A-C)** Labelling of the inputs to the colliculo-parabigeminal circuit. **A)** Injection strategy for labelling of the circuit connecting the retina to the parabigeminal nucleus, via the superior colliculus. **B)** Example retina with labelled ganglion cells innervating the colliculo-parabigeminal circuit. Scale bar = 500 μm. **C)** Zoomed-in version of C. Scale bar = 50 μm. **D)** 8 example retinal ganglion cells from either injection approach (parabigeminal nucleus or pulvinar). Left: en-face view of the dendritic tree. Right: side-view of the dendritic tree. Location of ChAT-bands is indicated with two grey lines. The cells have been broadly separated into four stratification groups: bistratified (first column), below ChAT-bands (second column), between ChAT-bands (third column), and above ChAT-bands (last column). **E)** Distribution of dendritic tree diameter of retinal ganglion cells that are part of the colliculo-pulvinar (LP; orange) and the colliculo-parabigeminal nucleus (Pbg; green) circuit. **p < .01 Kolmogorov-Smirnov test. **F)** Retinal ganglion cell diameters for cells stratifying below, between, and above ChAT-bands. **G)** Retinal ganglion cells of each circuit were grouped into four stratification groups based on the peak of their dendritic profile. **p < .01 Kolmogorov-Smirnov test and Wilcoxon rank sum test. See also Figure S1.

The anatomy of labelled ganglion cells was recovered by staining the retinas with antibodies against GFP (binding to the GCaMP6s) and ChAT, an internal marker of depth formed by starburst amacrine cells (Sanes and Masland, 2014; Sümbül et al., 2014). We created high-resolution confocal image stacks of each ganglion cell (X, Y, Z: 0.38, 0.38, 0.25-0.35 μm/pixel). Applying a semi-automated image processing routine, we created a flattened version of each ganglion cell relative to the ChAT bands that enables a precise quantification of their dendritic morphology (Sumbul et al., 2014; Sümbül et al., 2014).

### Anatomy of retinal inputs to the colliculo-parabigeminal and the colliculo-pulvinar circuit

The morphology of 301 ganglion cells innervating the colliculo-parabigeminal (n = 146) and colliculo-pulvinar (n = 155) circuit were extracted. The cells showed a variety of morphologies: ~10% had bistratified dendritic trees (n = 31); ~20% were mono-stratified with dendrites below the ChAT-bands (n = 60); ~41% had their dendrites restricted to the region between the ChAT-bands (n = 124); and ~29% had dendrites stratifying exclusively above the ChAT-bands (n = 86; Figure 1D). We calculated for each cell the area covered by the dendrites and created a depth profile of the dendritic tree relative to the ChAT bands (Figure S1J). Our dataset contains cells with dendritic field diameters ranging from 90 to 420 μm (median: 206 μm), similar to the reported range of 80 to 530 μm (Badea and Nathans, 2004; Bae et al., 2018; Coombs et al., 2006; Kong et al., 2005; Sun et al., 2002).

Comparing the size and stratification of retinal ganglion cells innervating the colliculo-parabigeminal and colliculo-pulvinar circuits revealed two basic trends. First, cells innervating the colliculo-parabigeminal circuit had larger dendritic trees (median: 232 μm) than the cells innervating the colliculo-pulvinar circuit (median: 186 μm; p < .01, Kolmogorov-Smirnov test; Figure 1E). This was true at each stratification level (below ChAT-bands: 280 μm (parabigeminal) vs 183 μm (pulvinar); between ChAT-bands: 185 μm vs 130 μm; above ChAT-bands: 234 μm vs 170 μm; Figure 1F). Second, the stratification depth of cells innervating each circuit had distinct distributions. While the colliculo-pulvinar circuit showed strong bias for sampling from neurons stratifying between (50.3%) and above (32.9%) the ChAT-bands, the colliculo-parabigeminal circuit sampled more evenly from each stratification level (bistratified 18.5%, below ChAT-bands 26.0%, between 31.5%, above 24.0%; Figure 1G). We found that these differences are not due to a bias in the retinotopic location of the sampled cells (Figure S1K-M).

### Retinal inputs to the parabigeminal and the pulvinar circuit differ in molecular signature

To determine if the observed anatomical differences are reflected in the identity of the retinal ganglion cells, we performed histological staining against molecular markers of ganglion cell types. First, the four alpha cell types were labelled using SMI32-antibody against neurofilament (Bleckert et al., 2014; Coombs et al., 2006; Huberman et al., 2008; Krieger et al., 2017; Peichl et al., 1987). We found that around half of all rabies-labelled cells innervating the two circuits are alpha-cells (colliculo-parabigeminal median: 42%, n = 3 retinas; colliculo-pulvinar median: 53%, n = 4 retinas; Figure 2). The four classes of alpha-cells in mice can be distinguished based on stratification depth: sustained ON-alpha cells have dendrites below the ChAT-bands; the transient ON- and transient OFF-alpha cells have dendrites between the ChAT-bands, and the sustained OFF-alpha cell has dendrites above the ChAT-bands (Krieger et al., 2017). To identify which of the four alpha cell types are part of each circuit, we acquired local z-stacks of SMI32^+^/GCaMP6s^+^ double labelled neurons (n = 91 cells in 3 mice for the colliculo-parabigeminal circuit; n = 90 cells in 3 mice for the colliculo-pulvinar circuit). Each neuron was manually classified based on dendritic stratification depth (Figure 2C and D). Both circuits sample from sustained and transient OFF-alpha cells (parabigeminal vs pulvinar median: 13% vs 20% sustained; 32% vs 29% transient OFF-cells; 100% corresponds to all GFP+ cells). In contrast, transient ON-cells mostly innervate the colliculo-pulvinar circuit (parabigeminal vs pulvinar median: 4% vs 17%), while sustained ON-cells are almost exclusively labelled in the parabigeminal experiments (parabigeminal vs pulvinar median: 10% vs 0%).

**Figure 2:**
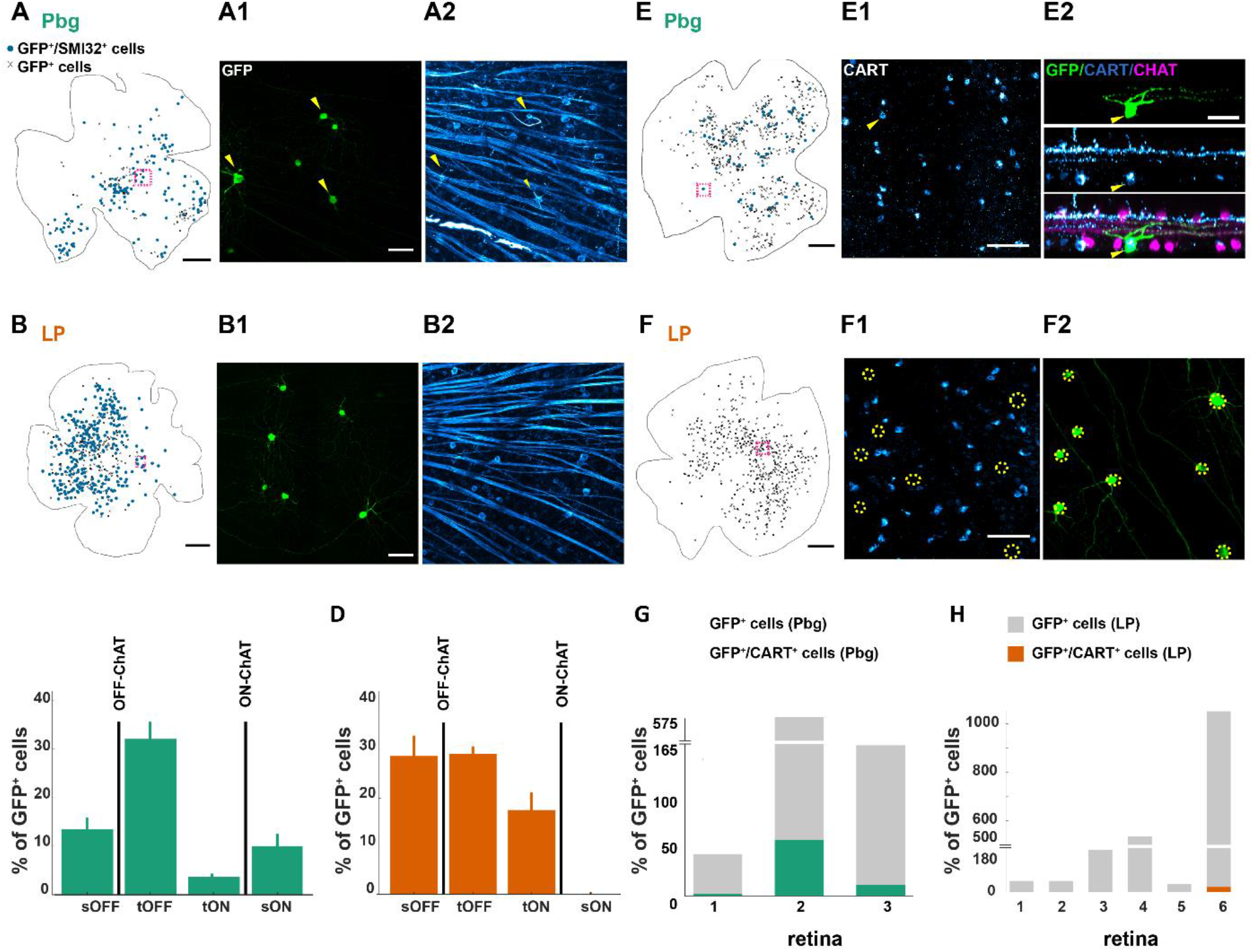
Distinct projection patterns of molecularly labelled retinal ganglion cells. **A-D)** Alpha retinal ganglion cells sending information to the colliculo-parabigeminal circuit. **A-B)** Example retina with SMI32-positive labelled retinal ganglion cells and examples of SMI32 positive neurons innervating the colliculo-parabigeminal and colliculo-pulvinar circuit, respectively. **A)** Example retina with SMI32-positive labelled retinal ganglion cells (blue dots) and SMI32-negative labelled retinal ganglion cells (other ganglion cells labelled after parabigeminal injections; black crosses). **A1)** Histological staining against GCaMP6s. Yellow arrows indicate SMI32-positive cells. **A2)** SMI32 histological staining against neurofilament. **C-D)** Median percentage of the four different alpha ganglion cell types (100% corresponds to all GCaMP6s-expressing cells). Colliculo-parabigeminal circuit n = 91 cells from 3 retinas. Bars indicate standard errors. Colliculo-pulvinar circuit n = 90 cells from 3 retinas. **E-H)** ON-OFF direction-selective cells labelled with CART. **E)** Example retina with CART-positive (dots) and CART-negative (crosses) labelled retinal ganglion cells. The arrow indicates a double-labelled cell (GFP signal not shown). **E2)** Side-view of the cell labelled in E1. The cell has been labelled by the rabies virus (GFP-positive; top) and is CART-positive (middle). Bottom: overlay of GFP, CART, and ChAT-staining. **F)** No CART^+^ neurons were labelled in pulvinar experiments. **G)** Number of CART^+^-cells in all 3 retinas of colliculo-parabigeminal circuit. **H)** Numbers of ON-OFF direction-selective cells are negligible in the colliculo-pulvinar circuit. Scale bar = 500 μm (A, B, E, F), scale bar = 50 μm (A1, A2, B1, B2, E1, E2, F1, F2).

In our dataset, the bistratified cells with dendritic density peaks aligned with the two ChAT-bands strongly resemble the morphology of ON-OFF direction-selective cells (Sanes and Masland, 2014). In the mouse retina, there are four types of ON-OFF direction-selective ganglion cells, each responding to one of the four cardinal directions. Three of the four types can be labelled with anti-CART antibodies (Dhande et al., 2013). We performed anti-CART histological staining in a subset of the retinas (Figure 2E-F). Double labelled neurons (GCaMP6s^+^ and CART^+^) are found almost exclusively after retrograde tracing from the parabigeminal nucleus (Figure 2G; median: 6.9% of all GCaMP6s-postive cells, range: 4.3 to 9.1%, n = 3 retinas). In the pulvinar experiments, a negligible percentage of the labelled ganglion cells are CART^+^ (Figure 2H; median: 1.3%, range: 0 to 2.1%, n = 6 retinas). In two of these retinas we saw no double labelled neurons (0/34 and 0/536).

### Clustering of ganglion cell anatomy reveals selective sampling by the colliculo-parabigeminal and the colliculo-pulvinar circuit

To get an estimate of the number of cell types innervating the colliculo-pulvinar and colliculo-parabigeminal circuits, we clustered our morphological data taking into consideration information about molecular identity (54/301 ganglion cells; n = 51 were SMI32^+^; n = 3 were CART^+^). We first set the average dendritic stratification profile for each molecularly identified cell type as a cluster centroid (4 SMI32^+^ types and 1 CART^+^ group). Then, all cells were clustered using affinity-propagation (Frey and Dueck, 2007). Our best estimate for the number of clusters was 12 based on three validation indices (Figure S3A-B). Dendritic tree location and size variance was consistent with previously published measurements, suggesting that our clustering result represented individual retinal ganglion cell types (Figure S3C-H). Within these 12 anatomical clusters, we found three distinct groups of cell types: 6 clusters that primarily innervate the parabigeminal circuit, 3 clusters that are almost exclusively part of the pulvinar circuits, and 3 clusters that innervate both circuits. Taken together, the colliculo-pulvinar and the colliculo-parabigeminal circuits sample from a small and only partially overlapping set of retinal ganglion cell types (Figure 3).

**Figure 3:**
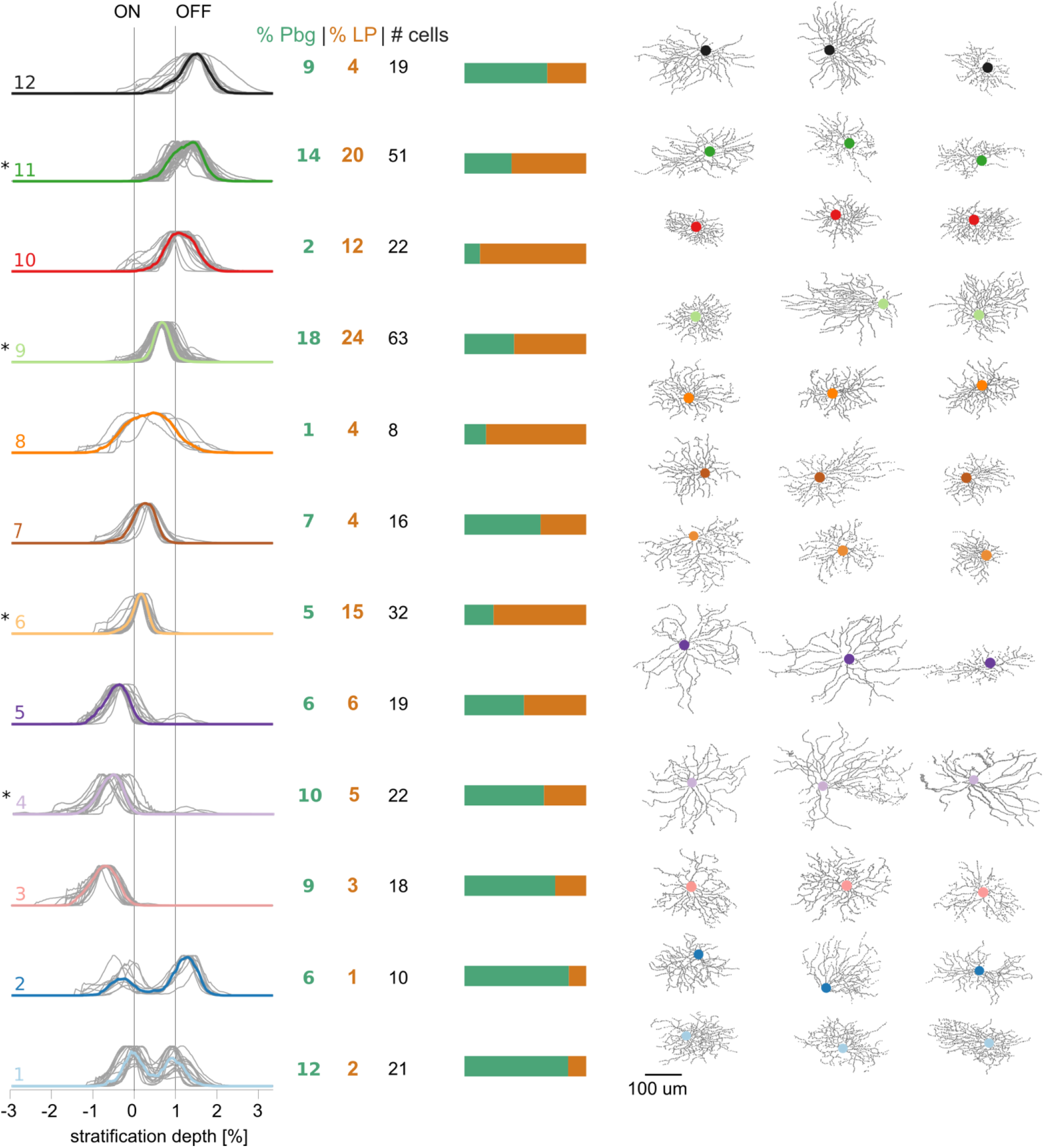
Retinal ganglion cell types targeting parabigeminal- and pulvinar-projecting collicular neurons. 12 clusters resulting from affinity propagation clustering based on the morphology and molecular markers of 301 retinal ganglion cells. Left: dendritic stratification profiles (median in colour). Profiles for individual cells are plotted in grey. Numbers indicate percentages of ganglion cells in a given cluster that were labelled after parabigeminal nucleus or pulvinar injections and the total number of cells in each cluster. Bars show proportion of cells in each cluster from parabigeminal (green) and pulvinar experiments (orange). * Indicates clusters with SMI32-positive cells. Right: en-face view of 3 example cells for each cluster that are the most similar to the cluster median. See also Figure S2.

The clusters specifically innervating the colliculo-parabigeminal circuit (cluster 1, 2, 3, 4, 7, and 12) include six putative types. These clusters included bistratified types: Cluster 1 contains the CART^+^ cells that have small dendritic fields with two dendritic layers that co-stratify with the ChAT-bands (n = 21; median diameter: 198 μm). These ganglion cells are the ON-OFF direction selective cell (Sanes and Masland, 2014). In cluster 2 we find a second bistratified cell type with a larger dendritic tree that stratifies below and above the ChAT-bands (n = 10; median diameter: 214 μm). Two clusters contain ON-neurons with dendrites below the ON-ChAT-band. One of the ON-types (cluster 4) contains SMI32^+^ cells (n = 22; median diameter: 276 μm), and has a morphology similar to sustained ON-alpha cells (Bleckert et al., 2014; Krieger et al., 2017). The very strong bias of this cluster for the colliculo-parabigeminal circuits mimics our antibody-labelling results of SMI32^+^ cells (Figure 2C and D). The second ON-type (cluster 3) does not contain SMI32^+^ cells, its dendrites lie further away from the ON-ChAT-band and it has a smaller dendritic tree (n = 18; median diameter: 232 μm). A single cluster (cluster 7) has neurons with dendrites between the ChAT-bands. These cells have relatively small dendritic trees that are located closer to the ON-ChAT band (n = 16; median diameter: 185 μm). The last cell type that specifically targets the colliculo-parabigeminal circuits consists of relatively large OFF-cells with a dendritic tree far above the OFF-ChAT-band (cluster 12, n = 19; median diameter: 243 μm).

The three ganglion cell types sending information preferentially to the colliculo-pulvinar circuit are in cluster 6, 8, and 10 and do not include any bistratified cells. The dendritic trees of the cells in cluster 6 are just above the ON-ChAT-band and this cluster contains SMI32^+^ neurons (n = 32; median diameter: 176 μm), which resemble the transient ON-alpha cell (Krieger et al., 2017). The selective routing of this type to the colliculo-pulvinar circuit is consistent with the molecular labelling experiments where transient ON-alpha cells are mostly labelled after pulvinar injections (Figure 2C and D). The cells in cluster 8 have an exceptionally broad dendritic tree and small dendritic fields (n = 8; median diameter: 199 μm). The third type of ganglion cells that uniquely targets the colliculo-pulvinar circuit is a rather small OFF-type with dendrites just above the OFF-ChAT-band (cluster 10, n = 22; median diameter: 166 μm).

Finally, three clusters innervate both circuits (cluster 5, 9 and 11). An ON-type (cluster 5) located just below the ON-ChAT-band has a sparse dendritic tree (n = 19; median diameter: 249 μm). Cluster 9 contains SMI32^+^ cells (n = 63; median diameter: 230 μm) with relatively large dendritic trees just below the OFF-ChAT-band that resemble the morphology of transient OFF-alpha cells (Huberman et al., 2008; Krieger et al., 2017; Münch et al., 2009). SMI32^+^ cells can also be found in cluster 11 (n = 51; median diameter: 187 μm). These OFF-cells have dendrites above the OFF-ChAT-band and are smaller in size, comparable to the published morphology of sustained OFF-cells (Bleckert et al., 2014; Farrow et al., 2013; Krieger et al., 2017). These results are consisted with our analysis of SMI32^+^ neurons (Figure 2C and D) where we found that both transient and sustained OFF-cells are labelled after parabigeminal and pulvinar injections.

### Functional classes of retina ganglion cells show refined pathway selectivity

We next characterized the functional response properties of the retinal ganglion cells innervating the colliculo-pulvinar and colliculo-parabigeminal circuits. To accomplish this, we performed two-photon targeted patch-clamp recordings from transsynaptically labelled neurons (Figure 4 and Figure S4). We presented each neuron with two sets of stimuli, one to determine its type (Figure 4) and a second to determine its responses to behaviourally relevant stimuli (Figure 7). We recorded from a total of 87 neurons, 47 innervating the colliculo-pulvinar circuit and 40 innervating the colliculo-parabigeminal circuit.

**Figure 4.**
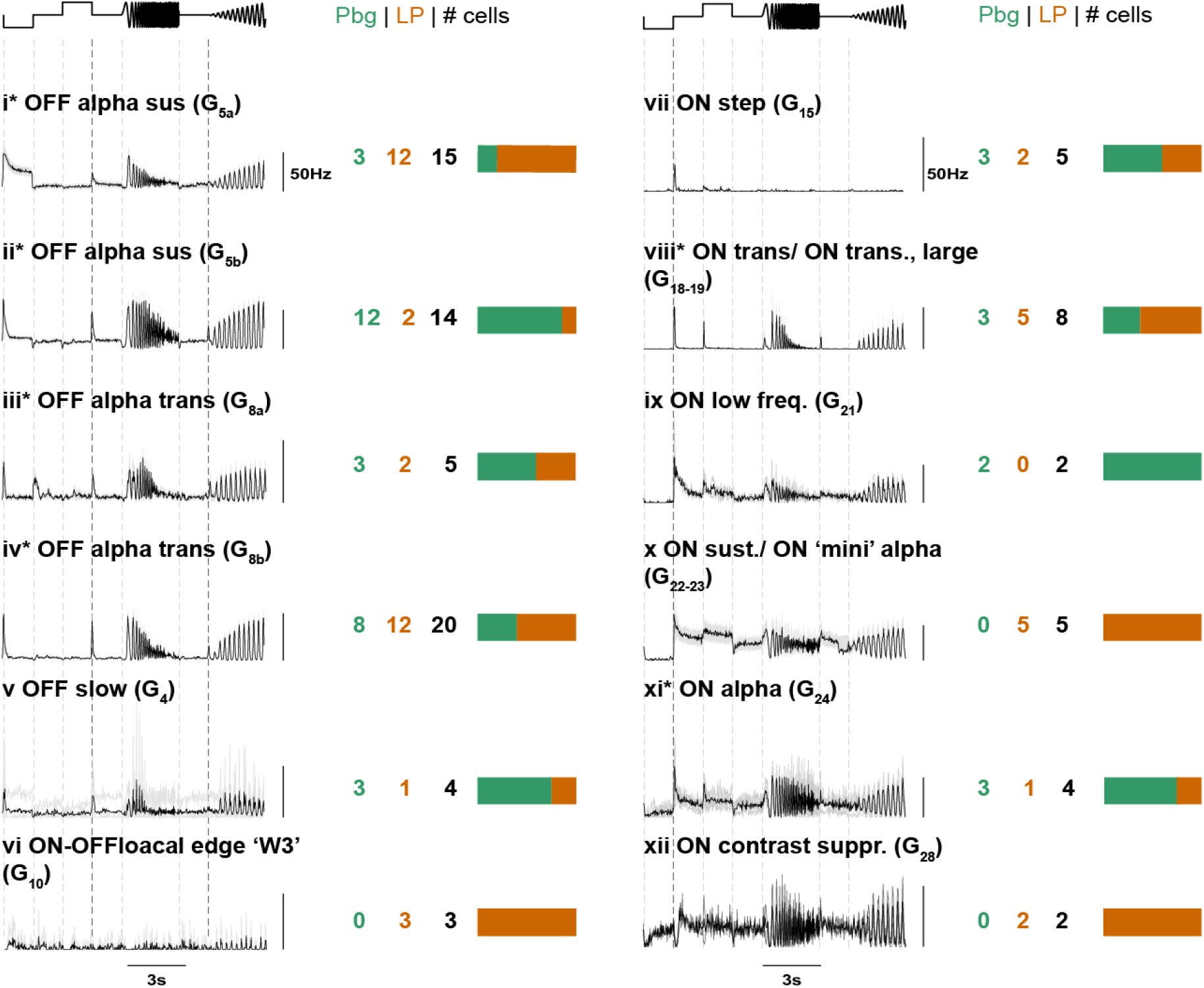
Functional response properties of labelled retinal ganglion cells. Average firing rate response (bin size 50 ms) of the labelled retinal ganglion cells to a full-field “chirp” stimulus. Mean response in black, standard error of the mean or the response of each cell in this group (if n<5) in grey. Numbers of recorded cells per circuit and group are indicated. Bar plots show the proportion of cells from either parabigeminal nucleus or pulvinar experiments in each group. * Indicates groups that include SMI32-positive cells.

To determine the functional class of each recorded cell we identified the best match group based on its response to a full-field stimulus (“chirp”) that Baden et al., 2016 previously used to classify retinal ganglion cells (hence refered to as G_xx_, Baden et al., 2016). In addition, a moving bar was presented to estimate each neurons direction and orientation tuning (Figure S4). In our recordings we identified 12 functional groups, where each functional group is consistent with the predicted responses of one of the anatomical clusters shown in Figure 3 (see Table S1). The functional groups included all alpha ganglion cells. However, we found response properties that separated the OFF alpha neurons into separate groups. Specifically, sustained OFF and transient OFF alpha ganglion cells each show two distinct response characteristics to the “chirp” stimulus that suggest that these anatomical classes form separate functional groups. The sustained OFF alpha neurons (G_5_ in Baden et al. 2016) could be split into two clear functional classes equivalent to G_5a_ and G_5b_, group *i* and *ii* respectively (Figure 4). The transient OFF alpha neurons could also be separated into two groups equivalent to G_8a_ and G_8b_, group *iii* and *iv*, respectively. Based on SMI32 staining and cell body size, sustained ON alpha neurons (G_24_; group *x*; median cell body size 195 μm^2^) could be distinguished from other sustained ON cells (G_22/23_; group *xi;* median cell body size 133 μm^2^). In addition, we identified transient ON alpha neurons (G_18/19_; group *viii*), ON contrast supressed neurons (G_28_; group *xii*), ON low frequency neurons (G_21_; group *ix*), ON step neurons (G_15_; group *vii*), ON-OFF local-edge ‘W3’ (G_10/14_; group *vi*) and OFF slow neurons (G_4_; group *v*). No recordings were made from direction-selective neurons as expected from our positive antibody staining for CART and bistratified anatomy inline with the ChAT bands (Figure 2 and 3).

We next asked whether the functional classes of retinal ganglion cells selectively innervate the colliculo-pulvinar or colliculo-parabigeminal pathways. In general, our functional recordings confirmed the pathway specific selectivity we observed in the molecular and anatomical analysis (Figure 2 and 3). Specifically, sustained ON alpha ganglion cells innervate the colliculo-parabigeminal pathway (group *xi* and Cluster 4); transient OFF alpha cells innervate both pathways (group *iii and iv* and Cluster 9); and local edge detectors (group *vi* and Cluster 8) selectively innervate the colliculo-pulvinar circuit. The putative ON low frequency (group *x* and Cluster 3) and putative OFF slow cells (group v and Cluster 12) preferentially innervate the colliculo-parabigeminal circuit. In addition, our functional recordings provided a more refined point of view of the selectivity for one of the putative cell classes. Specifically, the sustained OFF alpha ganglion cells (group *i* and *ii*, and Cluster 11), showed clear pathway specificity based on their functional response characteristics, which was not observed in the cluster based on molecular and anatomical data. We found group *i* (G_5a_) and group *ii* (G_5b_) innervated the colliculo-pulvinar and colliculo-parabigeminal circuit, respectively.

### Visual response properties of neurons in the pulvinar and parabigeminal nucleus

To test if the selective sampling of retinal inputs was reflected in the visual responses of the collicular targets, we performed multichannel silicon probe recordings in awake, head-fixed mice (Figure 5). In each recording session, stereotaxic coordinates were used to target the parabigeminal nucleus or pulvinar, the location of the probe was confirmed using histology and fluorescent dyes (Figure 5B and Methods). We recorded the brain activity on 384 electrodes spanning ~3800 μm in depth during visual stimulation (Figure 5C) and extracted the spikes for single units (Figure 5D). Both brain nuclei responded reliably to a set of visual stimuli including big-fast and small-fast objects as well as expanding dots (Figure 5E). However, the percentages of responding units differed for the different stimuli between the parabigeminal nucleus and the pulvinar (Figure 5F).

**Figure 5:**
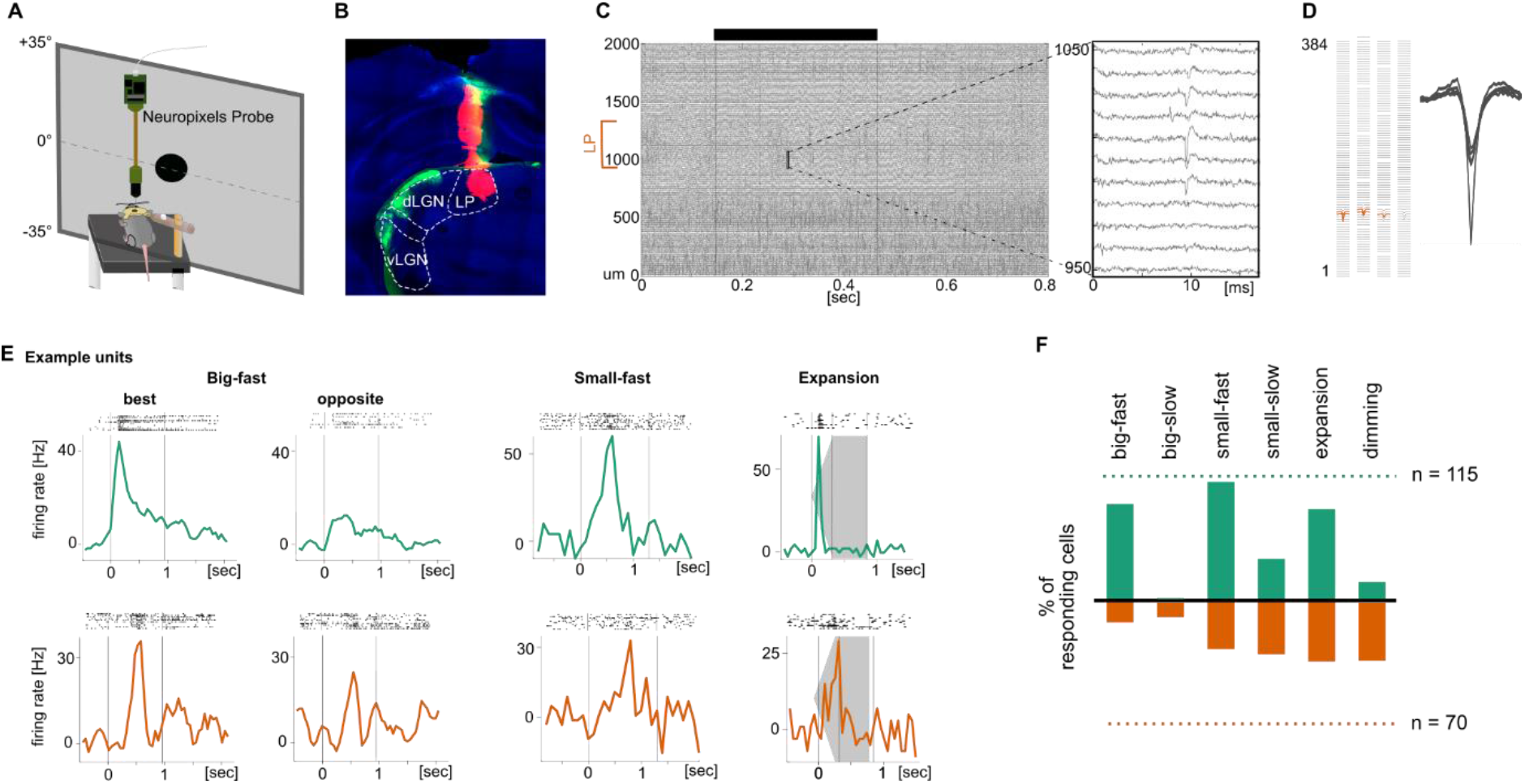
Recordings from the parabigeminal nucleus and pulvinar *in-vivo*. **A)** Scheme of setup for Neuropixels recordings in awake, head-fixed mice. **B)** Track of DiD-coated probe (magenta) visible in the pulvinar. Retina targets, including the LGN, were labelled using Choleratoxin-b-Alexa488 injections into the eye (green). **C)** Example raw trace on 201 out of 384 electrodes during the presentation of an expanding dot. The stimulus beginning and end are indicated with a black bar. The part of the probe covering the pulvinar is labelled. Right: 13 ms long snippet showing spiking activity on 10 electrodes. **D)** Foot print of a single neuron shown on the 384 electrodes and its 5 biggest template waveforms. **E)** Example responses from Pbg and pulvinar recordings to 10 repetitions of different stimuli. Stimuli were: Big-fast black square (53° side length, moving at 150°/sec); small-fast black dot (4° diameter, moving at 150°/sec); expanding black dot (expanded from 2° to 50° of diameter within 300 ms). The vertical lines indicate the stimulus beginning and end. **F)** Percentage of responding Pbg (green) and pulvinar (orange) units for four tested visual stimuli. 100% corresponds to the total number of light responsive units.

The responses of these two neuronal populations revealed a few key differences, and some similarities, in the visual response properties to biologically relevant stimuli. We found strong and reliable responses of neurons in the parabigeminal nucleus to the presentation of a fast-moving black square (53° side length, moving at 150°/sec) moving in 8 directions (Figure 5E). Most parabigeminal but very few pulvinar neurons responded to this stimulus (Figure 5F). The response amplitude and duration of responding neurons was similar for both nuclei (Figure S5). A large fraction of parabigeminal neurons showed a preference for one or two directions of motion (Figure 6A). Similar to the big-fast square, we found that most parabigeminal neurons responded with a sharp peak (median amplitude: 11 Hz, median half-width: 300 ms) to a small-fast dot (4° diameter, 150°/sec, Figure 6B). A large fraction of pulvinar neurons responded to this stimulus as well. Their responses were weaker but slightly sharper (median amplitude: 8 Hz, median half-width: 280 ms). Small-slow black dots (4° diameter, 21°/sec) generally induced weak, sluggish responses in rather few neurons in both brain regions (Figure 6C). However, pulvinar neurons tended to respond more sharply as indicated by a significantly smaller half-width of their response (pulvinar: 205 ms; Pbg: 505 ms). Finally, we tested responses to black-expanding dot (expanding from 2° to 50° of diameter within 300 ms) (Figure 6D). We found that both parabigeminal and pulvinar units responded strongly to the expanding dot. Responses to the disappearance of the black dot (return of screen to grey) were stronger in the pulvinar (median: 13 Hz, range: 2 to 78 Hz in pulvinar vs median: 10 Hz, range: 2 to 28 Hz in Pbg). Together, these findings suggest that the parabigeminal nucleus responds preferentially to movement of fast objects of different sizes moving in particular directions, while the pulvinar responds better to small and slower stimuli. Neurons in both nuclei are activated by expanding stimuli.

**Figure 6:**
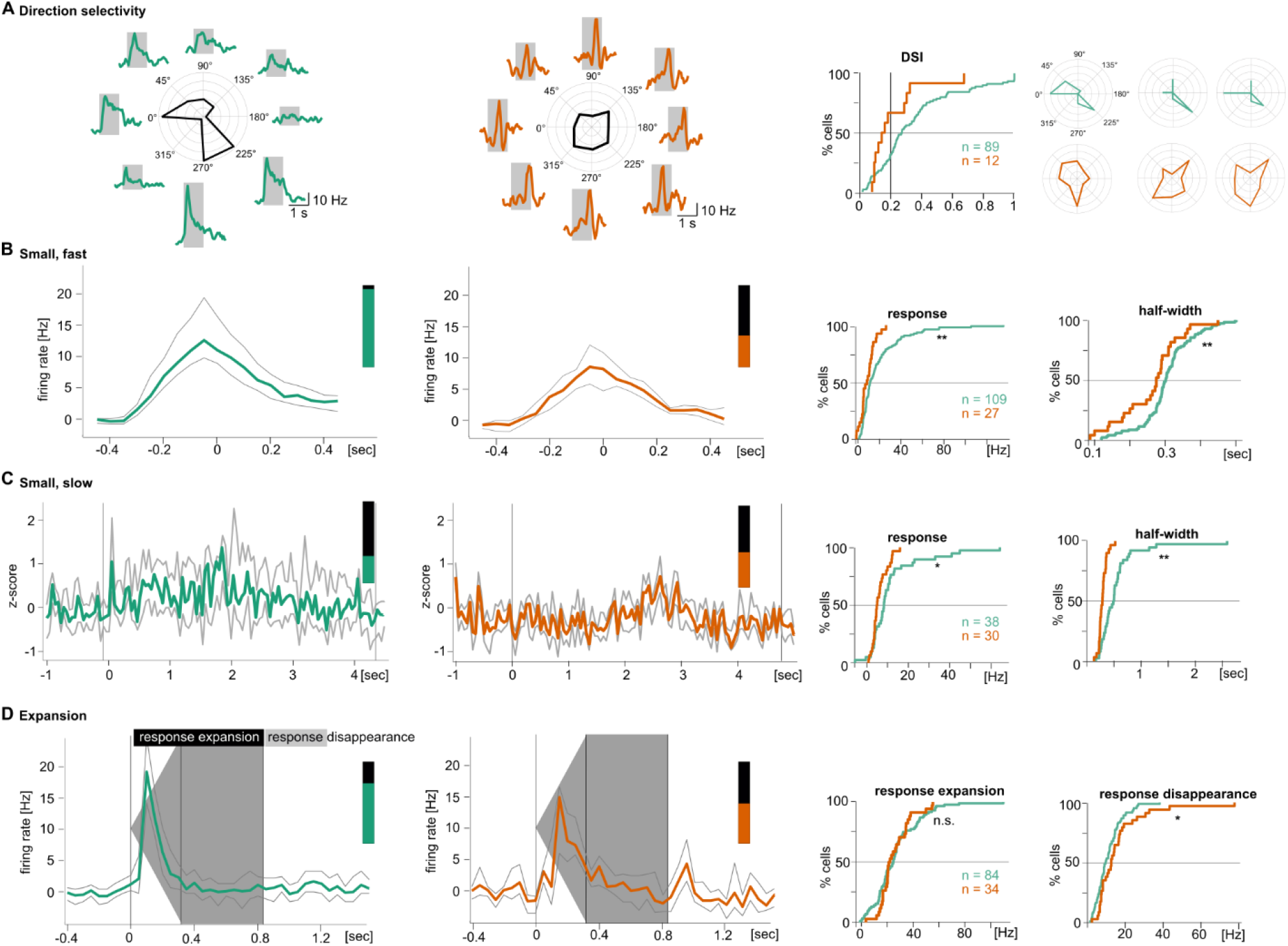
Neurons in the parabigeminal nucleus and pulvinar prefer different sets of visual stimuli. **A)** Direction selectivity was measured with a big-fast black square moving in 8 directions (53° side length, moving at 150°/sec). Left: Pbg example unit responding preferentially to a stimulus moving to the front and to stimuli moving to the back/down. Middle: Pulvinar example unit without direction preference. Right: Distribution of direction-selectivity indices (DSI) and three example cells with a DSI around the population average. **B)** Median ± octiles of peak response of all Pbg units (left; n = 109 out of 115 Pbg units with light responses) and all LP units (middle; n = 27 out of 70 LP units with light responses) to a small-fast dot (4° diameter, moving at 150°/sec). Right: cumulative distribution of maximal response amplitude and half-width. **C)** Median ± octiles of responses to a small-slow dot (4° diameter, moving at 21°/sec). Pbg: n = 38; LP: n = 30. Cumulative distributions as in B. **D)** Median ± octiles of responses to an expanding dot (from 2° to 50° of diameter within 300 ms). Pbg: n = 84; LP: n = 34. Cumulative distributions are shown for response amplitude during the expansion and for the response to the disappearance of the black dot. * p < .05, ** p < .01 Wilcoxon rank sum test.

### Differences in visual responses of pulvinar and parabigeminal nucleus explained by selective sampling of visual features

Our anatomical, molecular and physiological data shows that distinct visual information from the retina is wired to the parabigeminal nucleus and pulvinar. We therefore asked whether we could explain any of the observed response differences in the collicular targets by their selective sampling of visual information from the retina. We found two main differences in the response properties of the collicular targets. First, we observed a striking selectivity for motion direction in the parabigeminal nucleus that is absent in the pulvinar (Figure 6A). In accordance with these results, we found that direction-selective retinal ganglion cells exclusively innervated the colliculo-parabigeminal (Figure 2 and 3).

Second, parabigeminal and pulvinar neurons responded differently to a biologically salient stimulus consisting of an expanding black dot (Figure 6D). Over 80% of the parabigeminal neurons were strongly activated by the onset of the dot stimulus, out of which more than 3/4 responded exclusively to the onset (Figure S6 left). Pulvinar neurons rarely responded to the onset of the dot, but had more diverse activity patterns including responses to the expansion, the time after expansion when the dot was stationary at its final size, and to the disappearance of the dot (Figure S6 right). These different response patterns are reflected in the longer response latencies in the pulvinar population compared to the parabigeminal population (median latency 90 ms and mean latency 189 ms in pulvinar vs median latency 40 ms and mean latency 120 ms in parabigeminal nucleus) (Figure 7A). One of the major inputs to the colliculo-parabigeminal and colliculo-pulvinar circuits stems from sustained OFF alpha cells (cluster 11 in Figure 3), which could be separated into two functional groups *i* and *ii* that showed strong innervation preferences for the pulvinar and parabigeminal circuit, respectively (Figure 4). Interestingly, their responses to the expansion stimulus show comparable differences as their disynaptic targets (Figure 7B):

**Figure 7:**
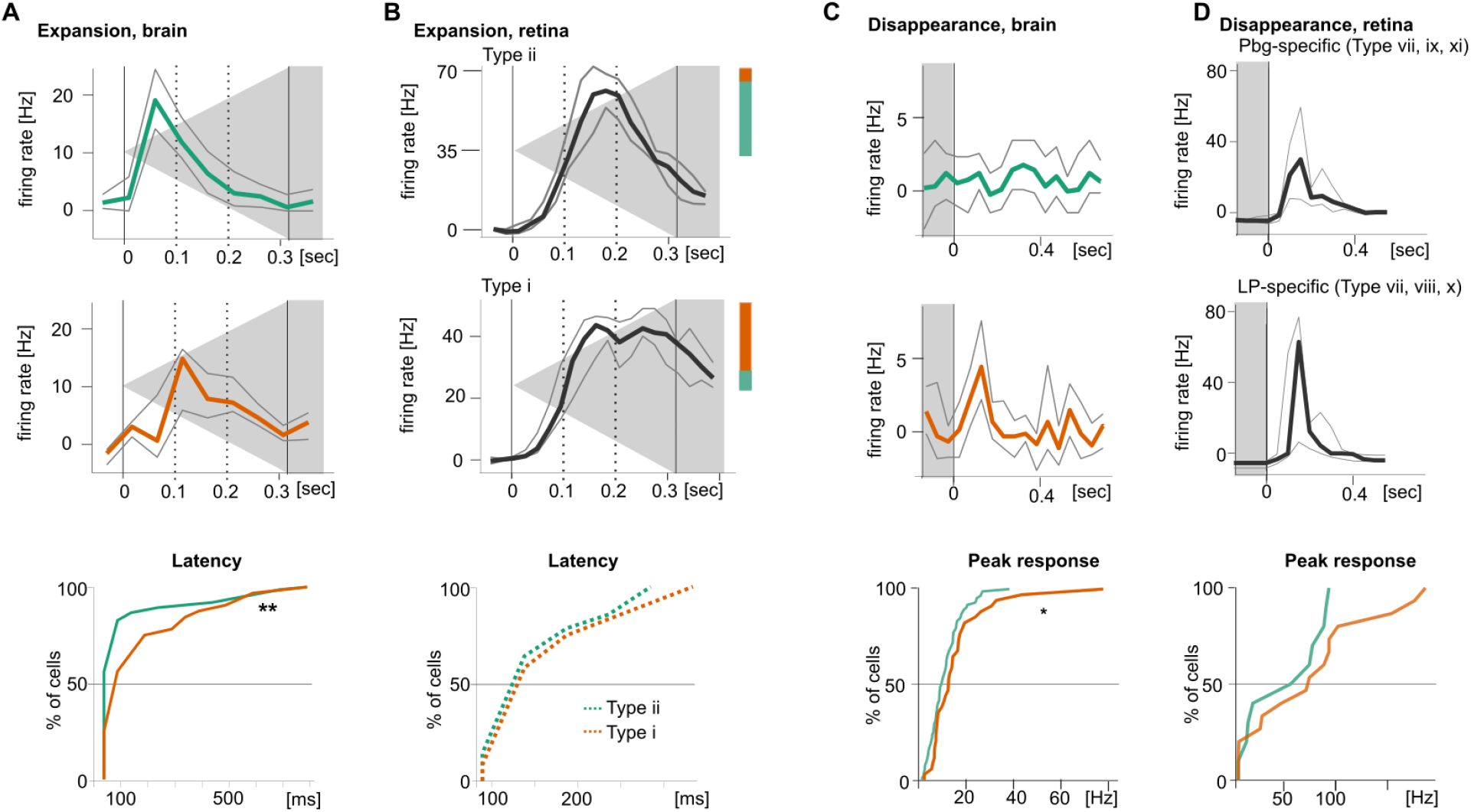
Differences in visual responses of pulvinar and parabigeminal nucleus explained by selective sampling of visual features. **A)** Population response to expanding dot stimulus for Pbg and pulvinar neurons. Bottom: Latency distributions of these responses. **B)** Population response to the same stimulus for Type *ii* and Type *i* ganglion cells. Coloured bars indicate percentages of ganglion cells from Pbg (green) and pulvinar experiments (orange). **C)** Population response to disappearance of expanding dot stimulus for Pbg and pulvinar neurons. Bottom: Response amplitude distributions. **D)** Population response to the same stimulus for ON-cells projecting to the parabigeminal (top) and pulvinar circuit (middle). * p < .05, ** p < .01 Wilcoxon rank sum test.

The parabigeminal-specific Type *ii* had a tendency towards faster responses than the pulvinar-specific Type *i* (median latency 125 ms and mean latency 154 ms in pulvinar vs median latency 125 ms and mean latency 167 ms in Type *ii*). Finally, the stronger response to the disappearance of the expanded dot in the pulvinar than in the parabigeminal nucleus (Figure 7C) is reflected in their respective retinal inputs where ganglion cells projecting to the pulvinar circuit show a stronger response than the parabigeminal-specific ones (median amplitude: 70 Hz in n = 15 pulvinar-specific cells vs 61 Hz in n =10 parabigeminal-specific cells) (Figure 7D).

## Discussion

By comparing the morphological, molecular and visual response properties of retinal ganglion cells innervating two different output pathways, parabigeminal nucleus and pulvinar, of the superior colliculus lead us to three conclusions (Figure 8). First, together the colliculo-parabigeminal and colliculo-pulvinar circuit sample from a limited set, ~14 out of more than 30 retinal ganglion cell types. Second, there is a clear segregation in the retinal ganglion cell types providing input to each circuit, where 7 putative ganglion cell types showed a strong preference for the colliculo-parabigeminal circuit, and 4 for the colliculo-pulvinar. Third, some of the response properties of neurons in the downstream targets could be explained by their selective sampling of different retinal ganglion cell types. These results support the notion that, in the superior colliculus, neural circuits are based on a dedicated set of connections between specific retinal inputs and different collicular output pathways.

**Figure 8.**
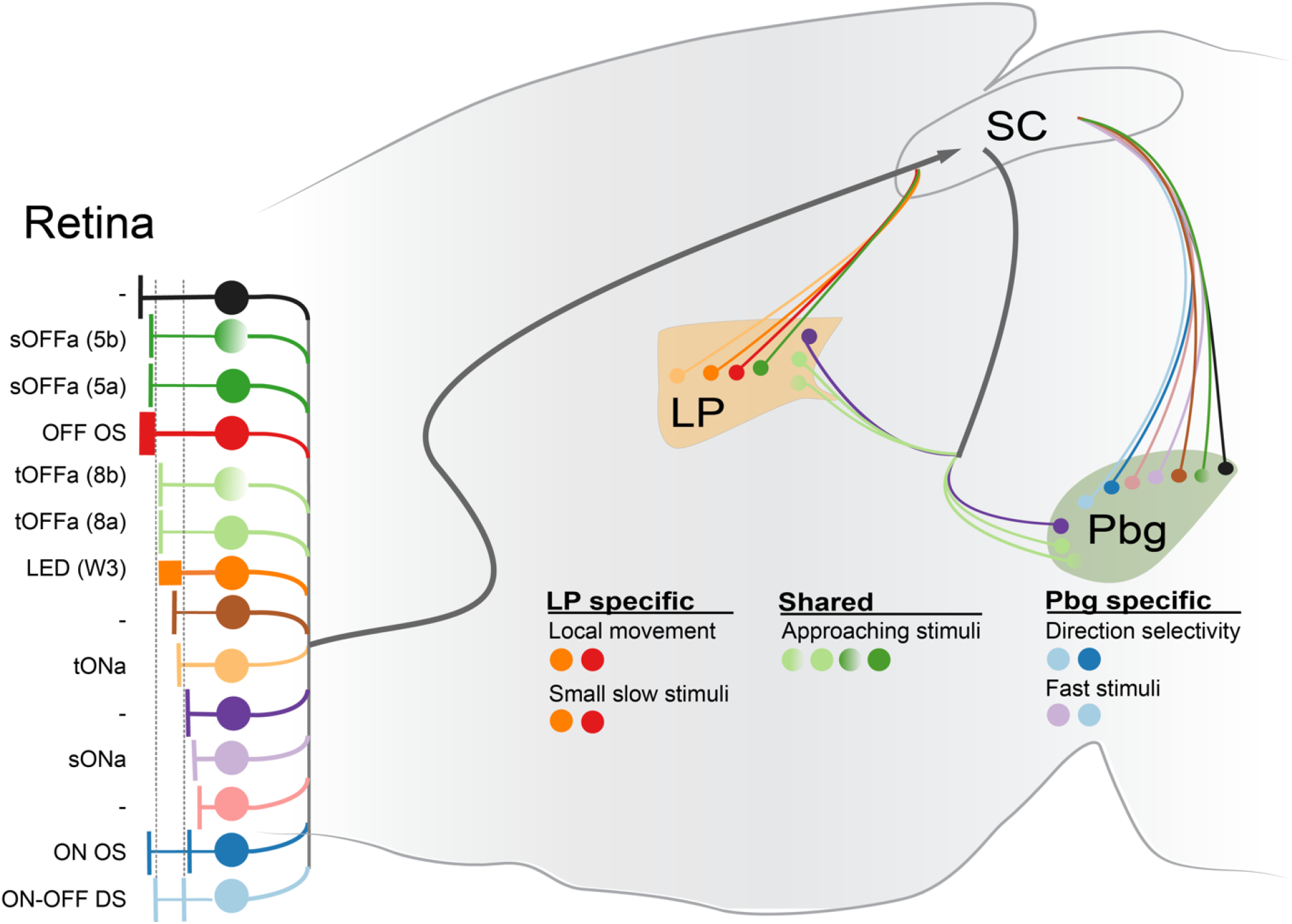
Schematic illustrating projection specific logic of sampling of retinal ganglion cell inputs to the superior colliculus. The color of the retinal ganglion cell types corresponds with the clusters defined in Figure 3. The functionally distinct sustained and transient OFF alpha ganglion cells (5a vs 5b and 8a vs 8b)) are indicated with shaded circles. The selective and shared response properties are shown with the putatively contributing retinal ganglion cell types indicated with coloured dots.

We used a combination of morphological, molecular, and physiological cues to link the 14 clusters to known retinal ganglion cell types (Table S1). Using these criteria, we could confidently identify 8 of the 14 putative retinal ganglion cell types, make a good prediction of 2 more, and limit the potential cell types for the other 4. The cells in cluster 1 are ON-OFF direction-selective cells, based on their characteristic dendritic structure that co-stratifies with the ChAT bands and positive CART labelling (Dhande et al., 2013; Sanes and Masland, 2014). The four alpha ganglion cell types (cluster 4, sustained ON alpha; Cluster 6 transient ON alpha, Cluster 9a/9b transient OFF alpha; Custer 11a/11b sustained OFF alpha) were positively identified based on their characteristic combination of positive SMI32 staining, dendritic anatomy and large cell body size (Bleckert et al., 2014; Krieger et al., 2017). Another cluster with indicative anatomical characteristics is cluster 8, containing cells that strongly resemble the “W3” cell, otherwise known as the “local-edge detector” or “object motion selective” cell (Farrow et al., 2013; Kim et al., 2010; Sümbül et al., 2014; Zhang et al., 2012). In addition, the second bistratified cluster 2 is most similar to ON orientation-selective neurons (Nath and Schwartz, 2016), and the small, dense OFF-cells in cluster 10 best match OFF orientation-selective neurons (Nath and Schwartz, 2017). There are three additional clusters that resemble cell types described by Sümbül and colleagues in terms of stratification pattern, size and dendritic density (Sümbül et al., 2014). First, the relatively dense ON-cells in cluster 3 have a similar morphology as the Kb-cells and fit best our physiological Type *ix* – putative ON low frequency cells. Second, the larger OFF-cells in cluster 12 are similar to Z-cells and might correspond to the slow OFF cells (Type v). Cluster 5 cells are large ON-cells with a sparse dendritic tree. They resemble the cells provisionally labelled as "U" by Sümbül and colleagues and the G10-cell by Völgyi and colleagues (Sümbül et al., 2014; Volgyi et al., 2009). Based on chirp response and morphology, it is unclear whether these cells correspond to our functional Type *vii* (ON step), *x* (ON sustained) or *xii* (ON contrast suppressed). Finally, the cells in cluster 7 stratifying just above the ON-ChAT-band show a similar profile as G5-cells (Volgyi et al., 2009).

The two neural circuits investigated here are each known to mediate visually guided aversive behaviours (Shang et al., 2015; Wei et al., 2015). In this context, the responses to biologically relevant stimuli of the ganglion cells innervating the two circuits are of interest. We found that neurons in the pulvinar respond poorly to large stimuli, but have sharper responses to small, slowly moving stimuli, which have been suggested to mimic a distant predator (Zhang et al., 2012). In addition, neurons in the pulvinar and parabigeminal nucleus respond well to quickly expanding dark stimuli, which are thought to mimic a quickly approaching threat (De Franceschi et al., 2016; Wei et al., 2015; Zhang et al., 2012). While robust responses to approaching stimuli have been reported in both pulvinar-projecting and parabigeminal-projecting collicular neurons, only pulvinar-projecting collicular neurons have been reported to respond to small slowly moving stimuli (Gale and Murphy, 2014, 2016, Shang et al., 2015, 2018). Consistent with these results, we found that the putative ganglion cell types providing selective input to the colliculo-pulvinar circuit have smaller receptive fields and two of them resemble cell types known to respond well to local motion (Cluster 6 and 8). In addition, transient OFF-alpha cells (Type *iii* and *iv* and Cluster 9) are known to preferentially respond to expanding stimuli and could mediate these responses in both circuits (Shang et al., 2015, 2018; Wei et al., 2015). Colliculo-parabigeminal specific retinal ganglion cells have larger dendritic fields and their putative function is to respond to large moving objects and their motion direction (Cluster 1, 2 and 4), and we found similar stimulus preferences in the parabigeminal neurons. Together these ganglion cells might detect predators attacking from angles that are not recognized by expansion-detectors.

While some of the differences between the response properties of the parabigeminal nucleus and pulvinar can be explained by their distinct retinal inputs, we recorded visual responses in the retinal ganglion cells that could not be detected in their disynaptic targets. For instance, we classified ganglion cells based on their responses to a full-field chirp stimulus, which did not evoke any response in neurons of the pulvinar or parabigeminal nucleus (Figure S5B). In addition, the colliculo-pulvinar circuit receives inputs from ganglion cells that respond well to big and fast objects, but responses to such stimuli were weak or absent in the pulvinar neurons. Shunting of these visual responses are likely mediated by local inhibitory circuitry in the colliculus, where removal of local inhibition has been demonstrated to reveal responses to large, stationary objects in pulvinar-projecting neurons (Gale and Murphy, 2016).

Studies investigating the organization of retinal inputs to single cells in the lateral geniculate nucleus have suggested that there is a large degree of fuzziness/variability in the information each neuron receives from the retina (Hammer et al., 2015; Liang et al., 2018; Morgan et al., 2016; Rompani et al., 2017). Here we demonstrate that in the superior colliculus a high degree of regularity exists if one considers the projection targets. These differences could exist either because this study was performed in the superior colliculus, which might have a more “hard-wired” architecture; or because we focused on projection specific disynaptic circuits instead of comparing inputs to single neurons. When considering the layer specific targets of the lateral geniculate nucleus in the visual cortex Cruz-Martin et al. suggest that direction-selective neurons are preferentially sampled (Cruz-Martín et al., 2014). We propose that understanding the specific input structure to neurons and cell types with different projection profiles (Gale and Murphy, 2014; Han et al., 2017) will greatly enhance our ability to create mechanistic models of how information from the sensory periphery informs the triggering of behaviours and decision making.

## Acknowledgements

We thank Keisuke Yonehara for supplying the Ntsr1-GN209-Cre mice, as well as Norma Kühn and João Couto for reading the manuscript. Grants are as follows: Marie-Curie CIG (631909) and FWO Research Project (G094616N) to K.F. This project has received funding from the European Union’s Horizon 2020 research and innovation programme under the Marie Skłodowska-Curie grant agreement No 665501 to K.R. (12S7917N). C.L. is funded by the Chinese Scholarship Council.

## Contributions

K.R., C.L. and K.F. conceived and designed the experiments and wrote the manuscript. K.R. and C.L. performed experiments and analysed the data. Q.D. implemented automated method for ChAT band detection. E.B. performed experiments. S.H. established the viral production facility.

## STAR★ METHODS

### KEY RESOURCES TABLE

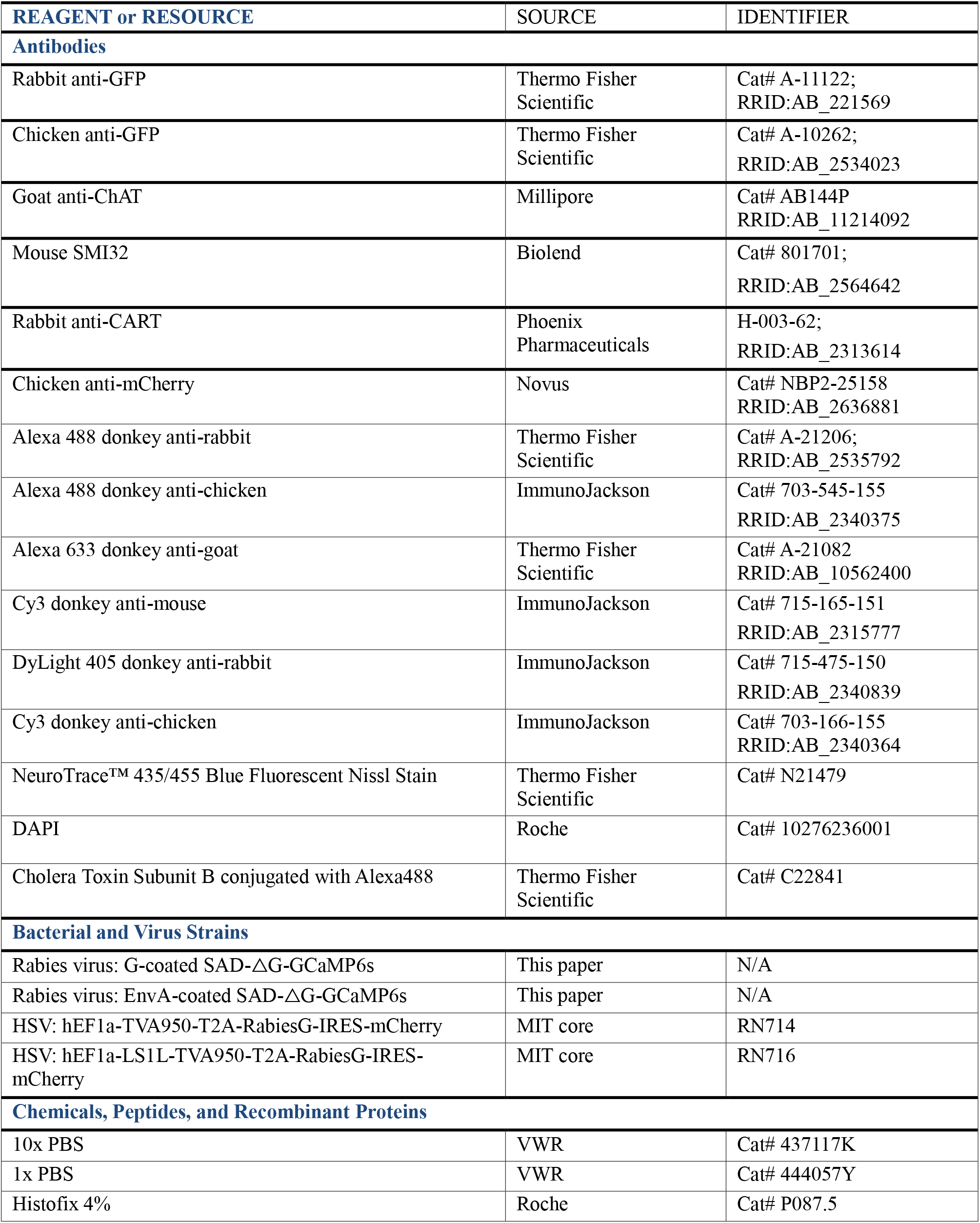
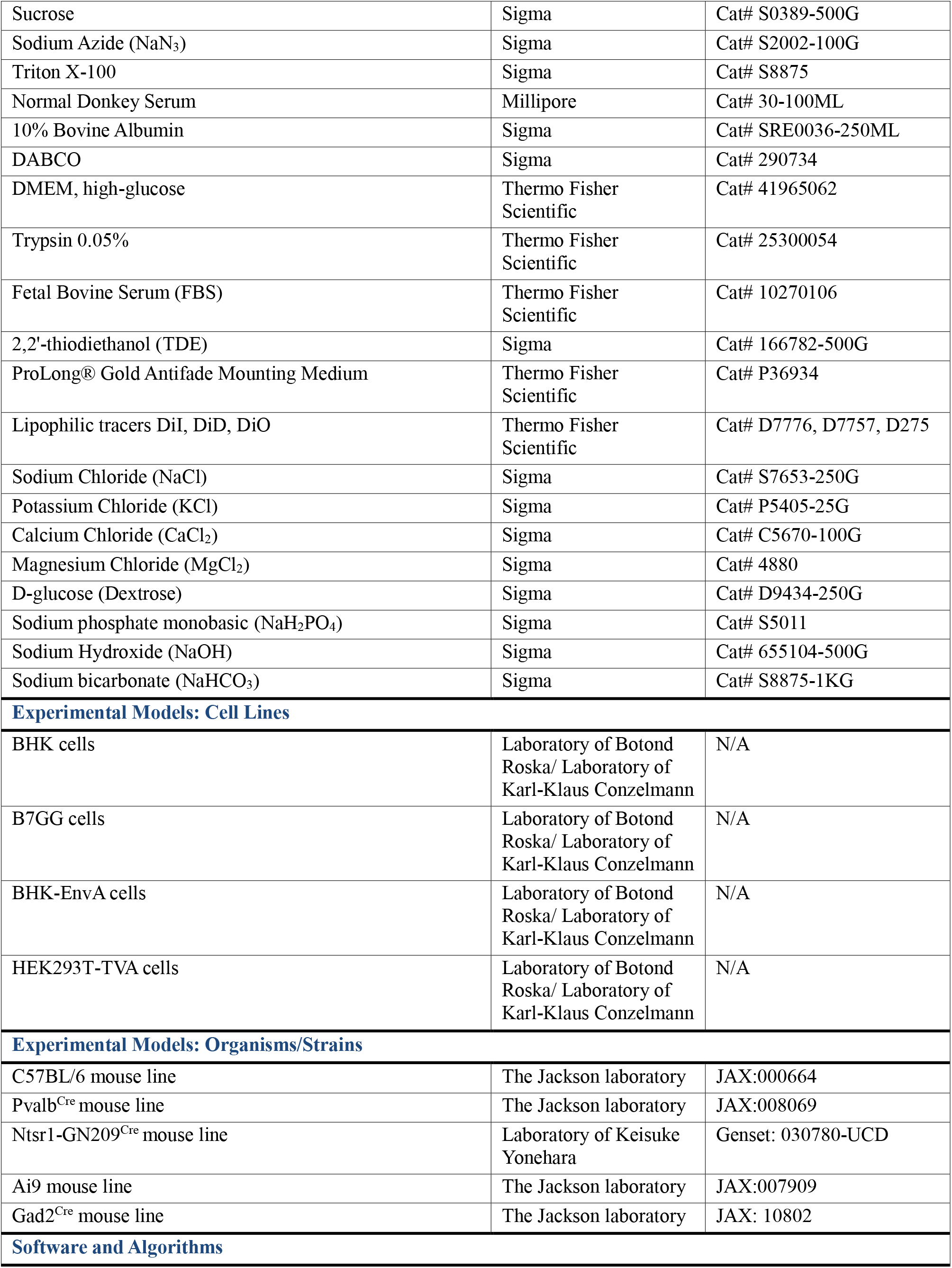
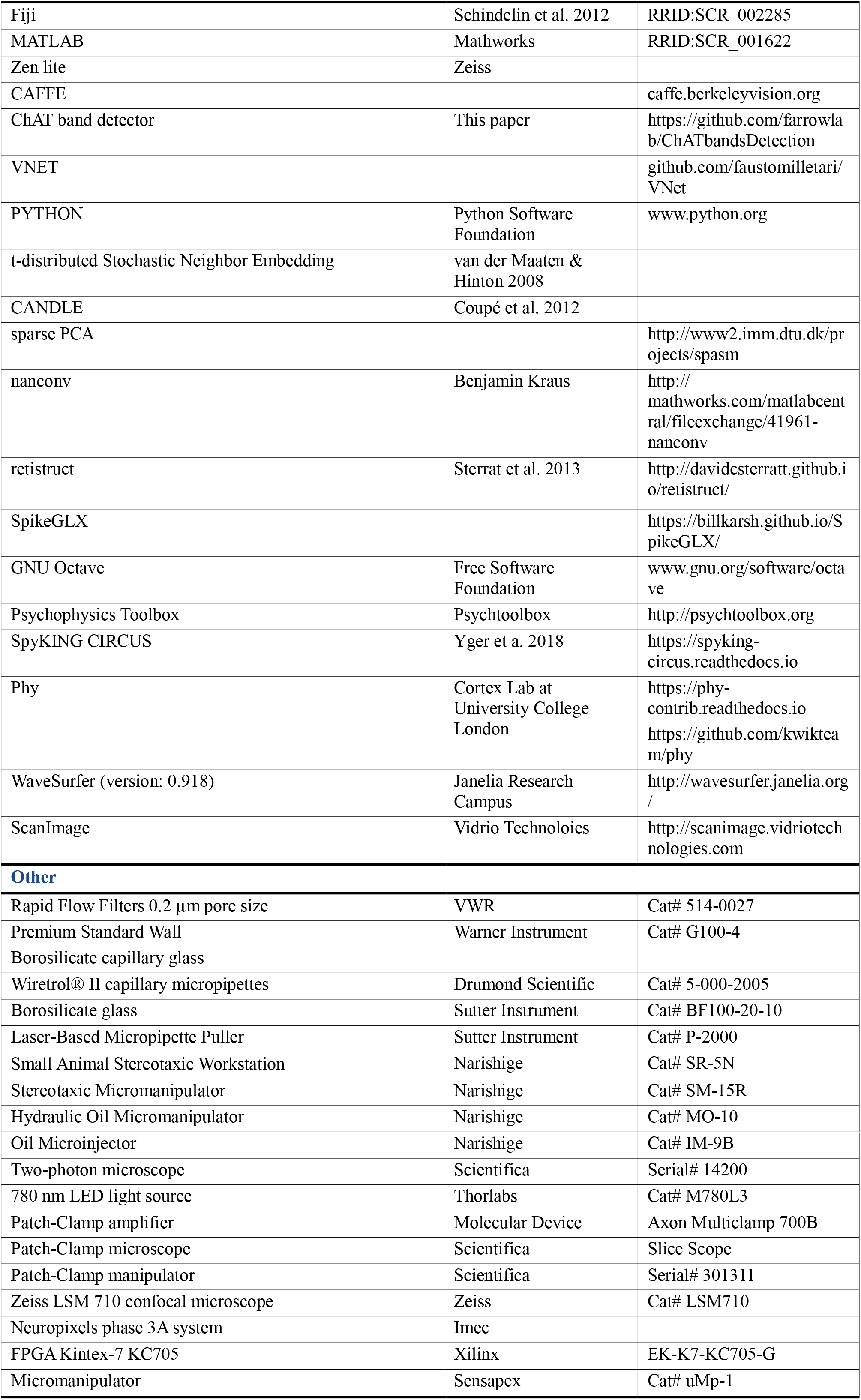

| REAGENT or RESOURCE | SOURCE | IDENTIFIER |
| --- | --- | --- |
| **Antibodies** |  |  |
| Rabbit anti-GFP | Thermo Fisher Scientific | Cat# A-11122;<br>RRID:AB_221569 |
| Chicken anti-GFP | Thermo Fisher Scientific | Cat# A-10262;<br>RRID:AB_2534023 |
| Goat anti-ChAT | Millipore | Cat# AB144P<br>RRID:AB_11214092 |
| Mouse SMI32 | Biolend | Cat# 801701;<br>RRID:AB_2564642 |
| Rabbit anti-CART | Phoenix Pharmaceuticals | H-003-62;<br>RRID:AB_2313614 |
| Chicken anti-mCherry | Novus | Cat# NBP2-25158<br>RRID:AB_2636881 |
| Alexa 488 donkey anti-rabbit | Thermo Fisher Scientific | Cat# A-21206;<br>RRID:AB_2535792 |
| Alexa 488 donkey anti-chicken | ImmunoJackson | Cat# 703-545-155<br>RRID:AB_2340375 |
| Alexa 633 donkey anti-goat | Thermo Fisher Scientific | Cat# A-21082<br>RRID:AB_10562400 |
| Cy3 donkey anti-mouse | ImmunoJackson | Cat# 715-165-151<br>RRID:AB_2315777 |
| DyLight 405 donkey anti-rabbit | ImmunoJackson | Cat# 715-475-150<br>RRID:AB_2340839 |
| Cy3 donkey anti-chicken | ImmunoJackson | Cat# 703-166-155<br>RRID:AB_2340364 |
| NeuroTrace™ 435/455 Blue Fluorescent Nissl Stain | Thermo Fisher Scientific | Cat# N21479 |
| DAPI | Roche | Cat# 10276236001 |
| Cholera Toxin Subunit B conjugated with Alexa488 | Thermo Fisher Scientific | Cat# C22841 |
| **Bacterial and Virus Strains** |  |  |
| Rabies virus: G-coated SAD-ΔG-GCaMP6s | This paper | N/A |
| Rabies virus: EnvA-coated SAD-ΔG-GCaMP6s | This paper | N/A |
| HSV: hEF1a-TVA950-T2A-RabiesG-IRES-mCherry | MIT core | RN714 |
| HSV: hEF1a-LS1L-TVA950-T2A-RabiesG-IRES-mCherry | MIT core | RN716 |
| **Chemicals, Peptides, and Recombinant Proteins** |  |  |
| 10x PBS | VWR | Cat# 437117K |
| 1x PBS | VWR | Cat# 444057Y |
| Histofix 4% | Roche | Cat# P087.5 |
| Sucrose | Sigma | Cat# S0389-500G |
| Sodium Azide (NaN<sub>3</sub>) | Sigma | Cat# S2002-100G |
| Triton X-100 | Sigma | Cat# S8875 |
| Normal Donkey Serum | Millipore | Cat# 30-100ML |
| 10% Bovine Albumin | Sigma | Cat# SRE0036-250ML |
| DABCO | Sigma | Cat# 290734 |
| DMEM, high-glucose | Thermo Fisher Scientific | Cat# 41965062 |
| Trypsin 0.05% | Thermo Fisher Scientific | Cat# 25300054 |
| Fetal Bovine Serum (FBS) | Thermo Fisher Scientific | Cat# 10270106 |
| 2,2'-thiodiethanol (TDE) | Sigma | Cat# 166782-500G |
| ProLong® Gold Antifade Mounting Medium | Thermo Fisher Scientific | Cat# P36934 |
| Lipophilic tracers DiI, DiD, DiO | Thermo Fisher Scientific | Cat# D7776, D7757, D275 |
| Sodium Chloride (NaCl) | Sigma | Cat# S7653-250G |
| Potassium Chloride (KCl) | Sigma | Cat# P5405-25G |
| Calcium Chloride (CaCl<sub>2</sub>) | Sigma | Cat# C5670-100G |
| Magnesium Chloride (MgCl<sub>2</sub>) | Sigma | Cat# 4880 |
| D-glucose (Dextrose) | Sigma | Cat# D9434-250G |
| Sodium phosphate monobasic (NaH<sub>2</sub>PO<sub>4</sub>) | Sigma | Cat# S5011 |
| Sodium Hydroxide (NaOH) | Sigma | Cat# 655104-500G |
| Sodium bicarbonate (NaHCO<sub>3</sub>) | Sigma | Cat# S8875-1KG |
| **Experimental Models: Cell Lines** |  |  |
| BHK cells | Laboratory of Botond Roska/ Laboratory of Karl-Klaus Conzelmann | N/A |
| B7GG cells | Laboratory of Botond Roska/ Laboratory of Karl-Klaus Conzelmann | N/A |
| BHK-EnvA cells | Laboratory of Botond Roska/ Laboratory of Karl-Klaus Conzelmann | N/A |
| HEK293T-TVA cells | Laboratory of Botond Roska/ Laboratory of Karl-Klaus Conzelmann | N/A |
| **Experimental Models: Organisms/Strains** |  |  |
| C57BL/6 mouse line | The Jackson laboratory | JAX:000664 |
| Pvalb<sup>Cre</sup> mouse line | The Jackson laboratory | JAX:008069 |
| Ntsr1-GN209<sup>Cre</sup> mouse line | Laboratory of Keisuke Yonehara | Genset: 030780-UCD |
| Ai9 mouse line | The Jackson laboratory | JAX:007909 |
| Gad2<sup>Cre</sup> mouse line | The Jackson laboratory | JAX: 10802 |
| **Software and Algorithms** |  |  |
| Fiji | Schindelin et al. 2012 | RRID:SCR_002285 |
| MATLAB | Mathworks | RRID:SCR_001622 |
| Zen lite | Zeiss |  |
| CAFFE |  | caffe.berkeleyvision.org |
| ChAT band detector | This paper | https://github.com/farrowlab/ChATbandsDetection |
| VNET |  | github.com/faustomilletari/VNet |
| PYTHON | Python Software Foundation | www.python.org |
| t-distributed Stochastic Neighbor Embedding | van der Maaten & Hinton 2008 |  |
| CANDLE | Coupé et al. 2012 |  |
| sparse PCA |  | http://www2.imm.dtu.dk/projects/spasm |
| nanconv | Benjamin Kraus | http://mathworks.com/matlabcentral/fileexchange/41961-nanconv |
| retistruct | Sterrat et al. 2013 | http://davidsterratt.github.io/retistruct/ |
| SpikeGLX |  | https://billkarsh.github.io/SpikeGLX/ |
| GNU Octave | Free Software Foundation | www.gnu.org/software/octave |
| Psychophysics Toolbox | Psychtoolbox | http://psychtoolbox.org |
| SpyKING CIRCUS | Yger et al. 2018 | https://spyking-circus.readthedocs.io |
| Phy | Cortex Lab at University College London | https://phy-contrib.readthedocs.io<br>https://github.com/kwikteam/phy |
| WaveSurfer (version: 0.918) | Janelia Research Campus | http://wavesurfer.janelia.org/ |
| ScanImage | Vidrio Technologies | http://scanimage.vidriotechnologies.com |
| **Other** |  |  |
| Rapid Flow Filters 0.2 µm pore size | VWR | Cat# 514-0027 |
| Premium Standard Wall Borosilicate capillary glass | Warner Instrument | Cat# G100-4 |
| Wiretrol® II capillary micropipettes | Drumond Scientific | Cat# 5-000-2005 |
| Borosilicate glass | Sutter Instrument | Cat# BF100-20-10 |
| Laser-Based Micropipette Puller | Sutter Instrument | Cat# P-2000 |
| Small Animal Stereotaxic Workstation | Narishige | Cat# SR-5N |
| Stereotaxic Micromanipulator | Narishige | Cat# SM-15R |
| Hydraulic Oil Micromanipulator | Narishige | Cat# MO-10 |
| Oil Microinjector | Narishige | Cat# IM-9B |
| Two-photon microscope | Scientifica | Serial# 14200 |
| 780 nm LED light source | Thorlabs | Cat# M780L3 |
| Patch-Clamp amplifier | Molecular Device | Axon Multiclamp 700B |
| Patch-Clamp microscope | Scientifica | Slice Scope |
| Patch-Clamp manipulator | Scientifica | Serial# 301311 |
| Zeiss LSM 710 confocal microscope | Zeiss | Cat# LSM710 |
| Neuropixels phase 3A system | Imec |  |
| FPGA Kintex-7 KC705 | Xilinx | EK-K7-KC705-G |
| Micromanipulator | Sensapex | Cat# uMp-1 |

## METHODS

### EXPERIMENTAL MODEL AND SUBJECT DETAILS

In total, 72 mice (3-5 weeks old for virus injections, 2-3 months for *in-vivo* physiology) of either sex were used in our experiments including PvalbCre, PvalbCre x Ai9, Ntsr1-GN209Cre, Ntsr1-GN209Cre x Ai9, and Gad2Cre. PvalbCre mice (JAX: 008069) (Hippenmeyer et al., 2005) express Cre recombinase in parvalbumin-expressing neurons. Ntsr1-GN209Cre mice (Genset: 030780-UCD) express Cre recombinase in Ntsr1-GN209-expressing neurons. Gad2Cre mice (JAX: 010802) express Cre recombinase in Gad2-expressing neurons. Ai9 (JAX: 007909) is a tdTomato reporter mouse line (Madisen et al., 2010). Animals were maintained on a 12-hour light/dark cycle, and fed with sterilized food, water, bedding and nesting material. All animal procedures were performed in accordance with standard ethical guidelines of KU Leuven and European Communities Guidelines on the Care and Use of Laboratory Animals (004-2014/EEC, 240-2013/EEC, 252-2015/EEC).

### METHOD DETAILS

#### Rabies virus production

Rabies production method was similar to previously published methods (Osakada and Callaway, 2013; Yonehara et al., 2013). Glycoprotein G-coated, G-deleted B19 rabies virus (G-coated SAD-ΔG-GCaMP6s RV) was amplified in B7GG cells, which express rabies glycoprotein G. For amplification, approximately 10^6^ infectious units of G-coated SAD-ΔG-GCaMP6s RV were used to infect five 10-cm plates of 80% confluent B7GG cells followed by 2-6 hours of incubation. Then, infected B7GG cells were treated with 0.05% trypsin and split into twenty-five 10-cm plates. To harvest the virus, we collected the supernatant of the infected cells every 3 days. 5-6 harvests were performed. To concentrate the virus, the supernatant was firstly centrifuged at 2,500 RPM and filtered (VWR, 514-0027) to get rid of the cell debris. Then the virus was spun in an ultracentrifuge for 5-12 hours at 25,000 RPM and at 4°C. After ultracentrifugation, the supernatant was discarded, and the pellet was dissolved in 200 μl of the original cell culture supernatant. The virus was tittered by counting a culture of infected BHK cells. To produce EnvA-coated SAD-ΔG-GCaMP6s RV, approximately 10^6^ infectious units of G-coated SAD-ΔG-GCaMP6s RV were used to infect BHK-EnvA cells. The same procedure as for the G-coated RV amplification was then applied. EnvA-coated SAD-ΔG-GCaMP6s RV was tittered by infection of HEK293T-TVA cells. The titer used for injection ranged from 10^7^ to 10^9^ infectious units/ml (IU/ml).

#### Surgical procedures

Animals were quickly anesthetized with Isoflurane (Iso-vet 1000mg/ml) and then injected with a mixture of Ketamine and Medetomidine (0.75 mL Ketamine (100 mg/mL) + 1 mL Medetomidine (1 mg/mL) + 8.2 mL Saline). Mice were placed in a stereotaxic workstation (Narishige, SR-5N). Dura tear (NOVARTIS, 288/28062-7) was applied to protect the eyes. To label the ganglion cells in the parabigeminal nucleus circuit, we performed the surgery on wild type mice and injected herpes-simplex-virus (HSV, hEF1a-TVA950-T2A-rabiesG-IRES-mCherry, MIT viral core, RN714) and EnvA-coated SAD-ΔG-GCaMP6s RV. In our experiment, we used PV-Cre mice as wild type mice. For the first injection of HSV into the parabigeminal nucleus, we used micropipettes (Wiretrol^®^ II capillary micropipettes, Drumond Scientific, 5-000-2005) with an open tip of 30 μm and an oil-based hydraulic micromanipulator MO-10 (Narishige) for stereotactic injections. Alternatively, we used an oil-based microinjector IM-9B (Narishige) with the corresponding micropipettes (Warner Instrument, G100-4) with an open tip of 30 μm. The injection coordinates for a 4 weeks old mouse with a bregma-lambda distance of 4.7 mm were AP: -4.20; ML: ±1.95; DV: 3.50 mm. As the mice were different in body size, we adjusted the coordinates for each mouse according to their bregma-lambda distance. To label the injection sites, DiD (Thermo, D7757) was used to coat the pipette tip. We injected in total 100-400 nl HSV in single doses of up to 200 nl with a waiting time of 5-10 min after each injection. Twenty-one days later, we injected rabies virus (EnvA-coated SAD-ΔG-GCaMP6s) into the superior colliculus using the same method as for the HSV injections. The retinotopic location of the first injection into the parabigeminal nucleus or the pulvinar is unknown. To maximize the labelling of ganglion cells in the retina, we thus covered as much as possible of the superficial layer of the superior colliculus during the second injection. We injected 100-200 nl of rabies virus at a depth of 1.7 – 1.8 mm at 4 different locations within a 1 mm^2^ field anterior of lambda and starting at the midline.

To label the pulvinar circuit, we performed the surgery on Ntsr1-GN209-Cre mice and injected a conditional HSV (hEF1a-LS1L-TVA950-T2A-RabiesG-IRES-mCherry, MIT viral core, RN716) and EnvA-coated SAD-ΔG-GCaMP6s RV. The injections into pulvinar and superior colliculus were the same as described for the parabigeminal nucleus. The injection coordinates for the pulvinar in a 4 weeks old mouse with a bregma-lambda distance of 4.7 mm were AP: -1.85; ML: ±1.50; DV: 2.50 mm.

Following injection, the wound was closed using Vetbond tissue adhesive (3M,1469). After surgery, mice were allowed to recover on top of a heating pad and were provided with soft food and water containing antibiotics (emdotrim, ecuphar, BE-V235523).

#### Retina Immunohistochemistry

Mouse retinas were extracted eight days after the rabies virus injection into the superior colliculus. After deep anaesthesia (120μl of Ketamine (100mg/ml) and Xylamine (2%) in saline per 20g body weight), eyes were gently touched with a soldering iron (Weller, BP650) to label the nasal part of the cornea and then enucleated. The retinas were extracted in 1x PBS (Diluted from 10x PBS (VWR, 437117K), pH 7.4) and three cuts were made to label the nasal, dorsal and ventral retina.

The dissected retinas were fixed in 4% paraformaldehyde (Histofix, ROTH, P087.5mm) with 100 mM sucrose for 30 min at 4°C, and then transferred to a 24-well plate filled with 1x PBS and washed 3 times for 10 min at room temperature or transferred into 15 ml 1x PBS and washed overnight or longer at 4 °C. After washing, retinas were transferred to wells containing 10% sucrose in 1x PBS with 0.1% NaN_3_ (w/v) and allowed to sink for a minimum of 30 min at room temperature. Then retinas were transferred to wells containing 20% sucrose in 1x PBS with 0.1% NaN_3_ (w/v) and allowed to sink for a minimum of 1 hour at room temperature. Finally, retinas were put into 30% sucrose in 1x PBS with 0.1% NaN_3_ (w/v) and allowed to sink overnight at 4°C. The next day, freeze-cracking was performed: retinas were frozen on a slide fully covered with 30% sucrose for 3-5 min on dry ice. The slides were then thawed at room temperature. The freeze-thaw cycle was repeated two times. Retinas were washed 3 times for 10 min each in 1x PBS, followed by incubation with blocking buffer (10% NDS, 1% BSA, 0.5% TritonX-100, 0.02% NaN_3_ in 1x PBS) for at least 1 hour at room temperature. Primary antibody solution was added after blocking and retinas were incubated for 5-7 days under constant gentle shaking at room temperature. Primary antibodies were rabbit anti-GFP (Invitrogen, A-11122, 1:500) and goat anti-ChAT (Chemicon, Ab144P, 1:200). They were prepared in 3% NDS, 1% BSA, 0.5% TritonX-100, 0.02% NaN_3_ in 1x PBS. After incubation, retinas were washed 3 times for 10 min in 1x PBS with 0.5% TritonX-100 before being transferred into the secondary antibody solution (Alexa488 donkey anti-rabbit (Invitrogen, A21206, 1/500) and Alexa633 donkey anti-goat (Invitrogen A-21082, 1:500); prepared in 3% NDS, 1% BSA, 0.5% TritonX-100, 0.02% NaN_3_ in 1x PBS). Nuclei were stained with DAPI (Roche, 10236276001, 1:500) together with the secondary antibody solution. The retinas were incubated in the secondary antibody with DAPI solution overnight at 4 °C. Retinas were then washed 3 times in 1x PBS with 0.5% TritonX-100 and 1 time in 1x PBS. Before mounting, the water in the sample was exchanged with different concentrations of 2,2’-Thiodiethanol (TDE) (Sigma, 166782-500G) buffer (10% -> 25% -> 50% -> 97%) (Staudt et al., 2007). Then the retinas were embedded in ProLong^®^ Gold Antifade Mountant (Thermo, P36934) and gently covered with a #0 coverslip (MARIENFEL, 0100032, No.0, 18*18 mm). To avoid squeezing the retinas, we put 4 strips of Parafilm (Parafilm, PM999) around the retina before adding the coverslip. Some of the retinas were mounted in 97% TDE with DABCO (Sigma, 290734) after immersion into TDE. Some retinas were mounted with ProLong^®^ Gold Antifade Mountant directly after washing. Afterwards, nail polish was used to prevent evaporation and the samples were stored in darkness at 4 °C.

#### Retina Immunohistochemistry (for RGC molecular marker staining)

Similar procedures were used to stain the retinas for neurofilament or CART. After fixation, freeze-cracking and blocking, primary antibody solution was added and the retinas were incubated for 5-7 days with gentle shaking at room temperature. Primary antibodies used were chicken anti-GFP (Invitrogen, A-10262, 1:500), goat anti-ChAT (Chemicon, Ab144P, 1:200), mouse SMI32 (Biolend, 801701,1:1000) and rabbit anti-CART (Phoenix, H-003-62,1/500). They were prepared in 3% NDS, 1% BSA, 0.5% TritonX-100, 0.02% NaN_3_ in 1x PBS. Retinas were washed 3 times, 15 min each, in 1x PBS with 0.5% TritonX-100 before being transferred into the secondary antibody solution consisting of Alexa488 donkey antichicken (ImmunoJackson, 703-545-155, 1:500) and Alexa633 donkey anti-goat (Invitrogen A-21082, 1:500), Cy3 donkey anti-mouse (ImmunoJackson, 715-165-151, 1:400) and DyLight™ 405 donkey antirabbit (ImmunoJackson, 715-475-150, 1:200) with 3% NDS, 1% BSA, 0.5% TritonX-100, 0.02% NaN_3_ in 1x PBS. Retinas were incubated in secondary antibody solution overnight at 4°C. Slices were washed 3 times for 10-15min each in 1x PBS with 0.5% TritonX-100 and 1 time in 1x PBS. After washing, the retinas were immersed in different concentrations of TDE buffer, then were mounted with either 97% TDE with DABCO or ProLong^®^ Gold Antifade Mountant. Some of the retinas were directly mounted with ProLong^®^ Gold Antifade Mountant without increasing concentrations of TDE.

#### Brain Immunohistochemistry

After removing the eyes, mice were immediately perfused with 1x PBS and 4% paraformaldehyde (PFA) and brains were post-fixed in 4% PFA overnight at 4 °C. Vibratome sections (100-200 μm) were collected in 1x PBS and were incubated in blocking buffer (1x PBS, 0.3% Triton X-100, 10% Donkey serum) at room temperature for 1 hour. Then slices were incubated with primary antibodies in blocking buffer overnight at 4 °C. The next day, slices were washed 3 times for 10 min each in 1x PBS with 0.3% TritonX-100 and incubated in secondary antibody solution diluted in blocking buffer for 2 hours at room temperature or overnight at 4 °C. Primary antibodies used were rabbit anti-GFP (Thermo Fisher, A-11122, 1:500) and chicken anti-mCherry (Novus, NBP2-25158, 1:1000) and secondary antibodies used were Alexa488 donkey anti-rabbit (Thermo Fisher, A21206, 1:500-1000) and Cy3 donkey anti-chicken (ImmunoJackson, 703-166-155, 1:800-1000). Nuclei were stained with DAPI (Roche, 10236276001, 1:500) together with the secondary antibody solution. Sections were then again washed 3 times for 10 min in 1x PBS with 0.3% TritonX-100 and 1 time in 1x PBS, covered with mounting medium (Dako, C0563) and a glass coverslip. For the Pbg experiments, we applied Nissl stain instead of the DAPI stain, where the Pbg can be identified as a cell-dense area. Nissl stain was applied after the secondary antibody staining. After washing with 1x PBS, the brain slices were incubated with Nissl in 1x PBS (NeuronTrace 435/455, Thermo, N21479, 1:150) for at least 20 minutes at room temperature. Afterwards, the sections were rinsed for 10 minutes in 1x PBS with 0.1% TritonX-100, followed by another 2 times washing for 5 minutes each in 1x PBS. Finally, the sections were washed on a shaker for 2 hours at room temperature or overnight at 4 °C in 1x PBS.

#### Confocal Microscopy

Confocal microscopy was performed on a Zeiss LSM 710 microscope. Overview images of the retina and brain were obtained with a 10x (plan-APOCHROMAT 0.45 NA, Zeiss) objective. The following settings were used: zoom 0.7, 4x4-tiles with 0 to 15% overlap, 2.37 μm/pixel resolution. For single retina ganglion cell scanning, we used a 63x (plan-APOCHROMAT 1.4 NA, Zeiss) objective. The following settings were used: zoom 0.7, 2x2-tiles or more (depending on size and number of cells) with 0 to 15% overlap. This resulted in an XY-resolution of 0.38 μm/pixel and a Z-resolution between 0.25 and 0.35 μm/pixel. The Z-stacks covered approximately 50 μm in depth.

#### *In-vivo* electrophysiology

##### Surgical procedure

8 PV-Cre mice of either sex at the age of 2-2.5 months were quickly anesthetized with Isoflurane (Iso-vet 1000 mg/ml) and then either maintained under Isoflurane anaesthesia or injected with a mixture of Ketamine and Medetomidine (0.75 mL Ketamine (100 mg/mL) + 1 mL Medetomidine (1 mg/mL) + 8.2 mL Saline). Lidocaine (0.5%, 0.007 mg/g body weight) was injected under the skin above the skull, the animal’s head was shaved, the skin and muscle tissue removed, and a titanium head plate fixed to the skull using dental cement (Metabond, Crown & Bridge). After recovery from anaesthesia animals were single-housed and were administrated Buprenorphine and Cefazolin for 60 hr post-surgery (Buprenorphine 0.2 mg/kg I.P. and Cefazolin 15 mg/kg I.P. in 12-hour intervals) and Dexamethasone (max. 0.2 ml of 0.1 mg/ml per day) depending on the condition of the animal. After this recovery phase animals were habituated for 3-4 days to the recording setup in sessions of increasing head-fixed time. One day before the first recording, the animals were anesthetized with Isoflurane and small craniotomies were performed (approximately 100 μm diameter, elongated to up to 300 μm laterally for parabigeminal coordinates and posteriorly for pulvinar coordinates). Coordinates were adjusted to each mouse’s skull size based on standard coordinates for a bregma-lambda distance of 4.7 mm. Standard coordinates pulvinar: bregma -2.0 / 1.7 lateral. Parabigeminal nucleus: bregma -4.2 / 2.0 lateral.

##### Data acquisition

Silicone Neuropixels probes phase 3A (Imec, Belgium) (Jun et al., 2017) were used to record light responses in the pulvinar and parabigeminal nucleus. The Neuropixels probes consist of a single shaft with 960 recording electrodes arranged in 480 rows with two electrodes each. The spacing between electrodes within a row (x) is 16 μm, and rows are 20 μm apart from each other (y) resulting in recording site length of 9600 μm. The 384 electrodes at the tip of the probe were recorded simultaneously in all experiments. Signals were split online into high-frequency (>300 Hz) and low-frequency (<300 Hz) and recorded separately at 30 kHz using the Neuropixels headstage (Imec), base-station (Imec) and a Kintex-7 KC705 FPGA (Xilinx). SpikeGLX was used to select recording electrodes, to calculate gain corrections and to observe and save the data. Stimulus timing information was recorded simultaneously using the digital ports of the base-station.

##### Presentation of Visual Stimuli

A calibrated 32-inch LCD monitor (Samsung S32E590C, 1920 x 1080 pixel resolution, 60 Hz refresh rate, average luminance of 2.6 cd/m2) was positioned 35 cm in front of the right eye, so that the screen was covering 100° of azimuth and 70° of altitude of the right visual field. Visual stimuli were presented on a gray background (50% luminance), controlled by Octave (GNU Octave) and Psychtoolbox (Kleiner et al., 2007). The following visual stimuli were used:

###### Large moving square

A black square of 53° side length moved with a speed of 150 °/sec across the screen in 8 direction (0°, 45°, 90°, 135°, 180°, 225°, 270°, 315°). Each direction was repeated 10 times.

###### Fast-small dot

A black dot of 4° diameter moved with 150 °/sec in two direction (left-right, right-left) at three different positions (centre, upper quarter, lower quarter) across the screen. Each position and direction was repeated 10 times.

###### Small-slow dot

Similar to the fast-small objects, a black dot of 4° diameter moved with 21 °/sec in two directions at three positions across the screen.

###### Expansion

A small dot linearly expanded from 2° to 50° of diameter within 300 ms at the centre of the screen. The stimulus was repeated 10 times.

###### Full-field “chirp” modulation

A full-field stimulus based on the “chirp” stimulus (Baden et al., 2016) starting with slow transitions gray-black-gray-white-gray (3 sec at each level), followed by a temporal modulation between black and white starting at 0.5 Hz and increasing to 8 Hz over a time of 6 sec. After 3 sec at a gray screen, the contrast was modulated from 0 to 100% over a time period of 5.5 sec at 2 Hz. The stimulus was repeated 10 times.

##### Experimental Design

Head-posted animals were fixed on a treadmill in front of the screen. For all pulvinar and some parabigeminal recordings, we coated the Neuropixels probe with a fluorescent dye (DiI, DiD or DiO, Thermo Fisher). The coordinates for the pulvinar (N = 4 recordings) or parabigeminal nucleus (N = 5) were measured again and the probe was slowly lowered into the brain using a micromanipulator. Some artificial cerebrospinal fluid (150 mM NaCl, 5 mM K, 10 mM D-glucose, 2 mM NaH_2_PO_4_, 2.5 mM CaCl_2_, 1 mM MgCl_2_, 10 mM HEPES adjusted to pH 7.4 with NaOH) was used to cover the skull. Then, the probe was lowered to the desired depth. In most cases, the probe was inserted further than the targeted brain area to ensure that the whole nucleus was covered. After 20-30 min, visual stimulation and recording of neural activity was started. The setup was covered with black curtains during the whole experiment.

##### Brain Histology for Probe Location

To facilitate the identification of the pulvinar and the correct location of the probe, we injected Cholera Toxin Subunit B conjugated with Alexa488 (Thermo Fisher) into the contralateral eye to label retinal targets such as the laterogeniculate nucleus of the thalamus. Then, the brain was fixed and Vibratome sections (100 μm) were collected in 1x PBS. The slices were washed in 1x PBS with 0.3% TritonX-100, then washed in 1x PBS and incubated for 20 min at RT with fluorescent Nissl Stain (NeuroTrace 435/455, Thermo Fisher, 1:150). Afterwards, the slices were washed in 1x PBS with 0.3% TritonX-100 and for at least 2h in 1x PBS. Brain slices were covered with mounting medium (Dako) and a glass coverslip, and imaged using a confocal microscope.

#### Retinal electrophysiology

##### Preparation of Retinas

For *in-vitro* recordings of retinal ganglion cells, we used mice that had been injected with herpes-simplex virus into the Pbg or pulvinar and rabies virus into the superior colliculus to label circuit specific retinal ganglion cells as described above. For pulvinar experiments, we analysed 47 cells from 14 Ntsr-Cre mice. For Pbg specific ganglion cells, we recorded 40 cells in retinas from PV-Cre (N = 8) or Gad2-Cre (N = 2) mice. Retinas were isolated from mice that were dark-adapted for a minimum of 30 minutes. Retina isolation was done under deep red illumination in Ringer’s medium (110 mM NaCl, 2.5 mM KCl, 1 mM CaCl_2_, 1.6 mM MgCl_2_, 10 mM D-glucose, 22 mM NaHCO_3_, bubbled with 5% CO_2_/95% O_2_, pH 7.4). The retinas were then mounted ganglion cell-side up on filter paper (Millipore, HAWP01300) that had a 3.5 mm wide rectangular aperture in the centre, and superfused with Ringer’s medium at 32–36°C in the microscope chamber for the duration of the experiment.

##### Electrophysiology

Electrophysiological recordings were made using an Axon Multiclamp 700B amplifier (Molecular Devices) and borosilicate glass electrodes (BF100-50-10, Sutter Instrument). Signals were digitized at 20 kHz (National Instruments) and acquired using WaverSurfer software (version: 0.918) written in MATLAB. The spiking responses were recorded using the patch clamp technique in loose cell-attached mode with electrodes pulled to 3-5 MΩ resistance and filled with Ringer’s medium. To visualize the pipette, Alexa 555 was added to the Ringer’s medium.

##### Targeted Recordings using Two-Photon Microscopy

Fluorescent cells were targeted for recording using a two-photon microscope (Scientifica) equipped with a Mai Tai HP two-photon laser (Spectra Physics) integrated into the electrophysiological setup. To facilitate targeting, two-photon fluorescent images were overlaid with the IR image acquired through a CCD camera. Infrared light was produced using the light from an LED. For some cells, z-stacks were acquired using ScanImage (Vidrio Technologies).

##### Presentation of Visual Stimuli

Stimuli were generated with an LCD projector (Samsung, SP F10M) at a refresh rate of 60 Hz, controlled with custom software written in Octave based on Psychtoolbox. The projector produced a light spectrum that ranged from ~430 nm to ~670 nm. The power produced by the projector was 240 mW/cm^2^ at the retina. Neutral density filters were used to control the stimulus intensity in logarithmic steps. Recordings were performed with filters decreasing the stimulus intensity by 1-2 log units. The following visual stimuli were used for retinal recordings:

###### Full-field “chirp” modulation

A full-field stimulus based on the “chirp” stimulus (Baden et al., 2016) starting with slow transitions gray-black-gray-white-gray (3 sec at each level), followed by a temporal modulation between black and white starting at 0.5 Hz and increasing to 8 Hz over a time of 6 sec. After 3 sec at a gray screen, the contrast was modulated from 0 to 100% over a time period of 5.5 sec at 2 Hz. The stimulus was repeated 10 times.

###### Spot-size

A black or white spot of 6 sizes (4°, 8°, 12°, 16°, 20°, 40°) was shown for 2 sec at the centre of the gray screen. Both the colours and the sizes were shown in random sequence.

###### Large moving bar

A black bar with a width of 40° moved with a speed of 150°/sec across the screen in 8 directions (0°, 45°, 90°, 135°, 180°, 225°, 270°, 315°). Each direction was repeated 5 times. The directions were randomized.

###### Expansion

A black dot linearly expanded from 2° to 50° of diameter within 300 ms (150°/sec) at the centre of the screen. The stimulus was repeated 10 times.

###### Dimming

A dot of 50° diameter linearly dimmed from background gray to black within 300 ms (150°/sec) at the centre of the screen. The stimulus was repeated 10 times.

###### Looming objects

A small dot non-linearly expanded from 2° to 50° of diameter at a slow (18.5°/sec), medium (92°/sec) and fast speed (150°/sec). Each condition was repeated 10 times.

###### Slow-small objects

A black dot of 4° diameter moved with 21 °/sec in two direction (left-right, right-left) at the centre line across the screen. Each direction was repeated 5 times.

##### Morphology of Patched cells

After patching, retinas were fixed and stained as described above. If the rabies labelling density allowed it, the morphology of the patched cells was imaged using a confocal microscope.

### QUANTIFICATION AND STATISTICAL ANALYSIS

#### Morphology of individual ganglion cells

The confocal Z-stacks were down-sampled and a threshold was applied to extract the dendritic tree. The position of the ChAT-planes was extracted and used to warp both the ChAT-signal as well as the binary Z-stack of the labelled cell. Then, dendrites from other cells, noise, and axons were removed and the position of the cell body was measured. The resulting warped dendritic tree was used for further analysis such as computation of the dendritic profile and area measurements. All code can be found on github (https://github.com/farrowlab/).

##### Down-sampling and binarization

The confocal Z-stacks of individual ganglion cells were denoised using the CANDLE package for MATLAB (Coupé et al., 2012) and down-sampled to have a resolution of XYZ = 0.5 x 0.5 x (0.25 to 0.35) μm per pixel and saved as MATLAB files. We then manually selected a threshold to transform the GFP-signal (i.e. the labelled cell) into a binary version where the whole dendritic tree was visible but noise was reduced as much as possible using an adapted version of the method described in (Sumbul et al., 2014; Sümbül et al., 2014).

##### Extraction of ChAT-positions

ChAT-band positions were either extracted manually or automatically using a convolutional neural network. For manual extraction, the ChAT-signal was smoothed using a two-dimensional standard-deviation filtering approach in the XY plane with a size of 21 x 21 pixels. The resulting Z-stacks were loaded into Fiji (Schindelin et al., 2012). ChAT-band positions were marked as described in (Sümbül et al., 2014). Briefly, we labelled points in the ON- and OFF-band with an approximate spacing of 20 μm in X- and Y-direction. For automated labelling, an end-to-end 3D Convolutional Neural Network called V-Net with a Dice Loss Layer (Milletari et al., 2016) was trained on noisy greyscale images of ChAT-images, to denoise and remove any cell bodies, creating a probability map of background and foreground, with foreground being voxels that might belong to the ChAT-bands. Two smoothness-regularized-least squares surfaces were fitted to manually labelled data to train the algorithm and to create ground truth binary masks. Then, Otsu’s thresholding method combined with connected component analysis was performed on the resulting probability map to automatically locate the points that belong to the ChAT-bands in new data-sets. Finally, two surfaces were independently fit to the corresponding data points to approximate the two ChAT-bands (https://github.com/farrowlab/ChATbandsDetection).

##### Warping

An adapted version of the code developed in the lab of Sebastian Seung was used to warp the GFP-signal (Sümbül et al., 2014). Briefly, the ChAT-band locations were used to create a surface map, which then was straightened in 3D-space. Then, the binarized GFP-signal was warped accordingly.

##### Soma position and removal of noise

After warping, the soma position was determined by filtering the GFP-signal with a circular kernel (adapted from (Sümbül et al., 2014)). If this method detected the soma, it was used to remove the soma from the GFP-data and the centre of mass was taken as the soma position. If this automated method failed, the soma position was marked manually. Afterwards, dendrites of other cells, axons, and noise were removed manually: The warped GFP-signal was plotted in side-view and en-face view in MATLAB and pixels belonging to the cell were selected manually.

##### Computation of the dendritic profile and area

The distribution of the cell’s dendritic tree was computed (Sümbül et al., 2014). Briefly, the Z-positions of all GFP-positive pixels were normalized to be between - 0.5 and 0.5. Then the Fourier transform of an interpolating low-pass filter was used to filter the Z-positions. This resulted in a vector containing the distribution of pixels in the Z-direction. If necessary, this profile was used to manually remove remaining axonal or somal pixels. In this case, the dendritic profile was computed again after cleaning of the data. The area of the dendritic tree was approximated by computing a convex hull (regionprops function in MATLAB). When diameters are given, they were calculated as D = 2*(area / π)^1/2^.

##### Down-sampling of dendritic tree for plotting

For en-face plots of the dendritic arbour, they were down-sampled by calculating the local neighbourhood median of all labelled pixels in patches of 50 x 50 pixels and with a sliding window of 10 pixels.

#### Clustering of retinal ganglion cell morphology

##### Affinity-propagation clustering

The dendritic stratification profiles of 301 ganglion cells were smoothed with the MATLAB function movmean (moving average with sliding window of 5 data points corresponding to 1.7 a.u. in stratification depth). The profiles of manually identified bistratified cells were set to negative values. The median dendritic profile of cells for which the molecular identity was known (SMI32 or CART), was calculated (4 different SMI32 cell types, 1 CART). Then the 5 first principle components of those medians and of the dendritic profiles of cells without known molecular identity were computed using sparse PCA (http://www2.imm.dtu.dk/projects/spasm/) and the similarity matrix of these principle components was calculated using the pdist function of MATLAB using Euclidean distance. Affinity-propagation (apcluster function in MATLAB) was used to cluster the similarity matrix with different preference values ranging from –1 to 0.6. The preference value for the 5 cluster centres based on SMI32- and CART-positive cells was always set to 1. Cells with known molecular identity were assigned to the clusters to whose median they contributed. Three validation indices (Calinski-Harabasz, Silhouette, Davies-Bouldin) were computed using the evalclusters function in MATLAB, normalized, and their median was used to determine the optimal preference value.

##### tSNE visualization

For visualization of the clustering result, we generated a five-dimensional non-linear embedding of the cells using t-distributed Stochastic Neighbor Embedding, tSNE (Van Der Maaten and Hinton, 2008). We used the smoothed dendritic profiles as input data, used cosine as a distance measurement, and set the number of PCA dimensions, which are calculated in a first step, to 25. For the graph in this paper, we show comparisons of the resulting tSNE dimension 1, 2, and 4.

#### Size distribution analysis

##### Retinal position

To calculate the visuotopic position of the analysed ganglion cells, we used the R-package retistruct (Sterratt et al., 2013), which morphs a flat retina with cuts onto a curvilinear surface. We manually labelled the nasal corner of the retina as well as tears and cuts. Reconstruction by retistrcut was then achieved by stitching the marked-up cuts; dividing the stitched outline into a mesh whose vertices then are mapped onto a curtailed sphere; and finally moving the vertices so as to minimise a physically-inspired deformation energy function (Sterratt et al., 2013). The resulting angle and radial distance from the optic nerve were used for plotting and quantification of retinal ganglion cell positions.

##### Comparison to PV cells

For size distribution comparisons, we used previously published data from 8 different types of parvalbumin-positive (PV) ganglion cells (Farrow et al., 2013). For each of our 12 clusters, we looked for a PV-type with a similar stratification depth and average dendritic field size. If there was such a PV-type, we calculated the median and quartiles of the dendritic field diameters of all cells of this type and compared it to our data.

Retinotopic size distribution: For retinotopic size distribution calculations, we computed a moving median diameter within a circular window of 250 μm radius, moving by 100 μm. The resulting 50 x 50 median size matrix was convolved with a gaussian with sigma = 200 μm (using MATLAB function fspecial and nanconv).

#### Quantification of SMI32+ cells and CART+ cells

##### Numbers of double-labelled cells

To quantify the number of double-positive cells for CART/GCaMP6s and SMI32/GCaMP6s, we scanned a z-stack (1 to 5 μm Z-resolution) of the whole retina using the confocal microscope with an 10x objective. Images of the anti-CART or SMI32 and the anti-GFP staining were opened in Fiji. For counting CART^+^ cells, cells were marked using the point tool and counted manually. Note that the anti-CART antibody also labels a group of amacrine cells, therefore the complete Z-stack should be checked for each CART^+^ cell to make sure that the labelling truly overlaps with the anti-GFP signal. The CART expression pattern was consistent with previous reports (Kay et al., 2011). In total we counted 3 retinas for parabigeminal experiments and 6 retinas for pulvinar experiments. For SMI32^+^ stainings, cells were counted manually using the cell counter plugin. In total we counted 3 retinas for parabigeminal experiments and 4 retinas for pulvinar experiments.

##### Numbers of cells for types of alpha cells

To test which of the four alpha cell types were part of each circuit, we acquired small high-resolution Z-stacks (2.5 μm/pixel) of XY = 103 x 103 μm size (128 x 128 pixel, 63x objective) covering the full depth of the dendritic tree and centred around the soma of 91 SMI32^+^ / GCaMP6s^+^ cells in n = 3 retinas from parabigeminal experiments and 90 SMI32^+^ / GCaMP6s^+^ cells in n = 3 retinas from pulvinar experiments. We plotted top and side views of each Z-stack in MATLAB and manually decided for each cell if it was a sustained ON-alpha cell (dendrites below the ON-ChAT band), a transient ON-alpha (dendrites just above the ON-ChAT band), a transient OFF-alpha (dendrites just below or on the OFF-ChAT band) or a sustained OFF-alpha cell (dendrites above the OFF-ChAT band).

#### Spike sorting

The high-pass filtered in-vivo data was automatically sorted into individual units using SpyKING CIRCUS (Yger et al., 2018). The following parameters were used: cc_merge = 0.95 (merging if crosscorrelation similarity > 0.95), spike_thresh = 6.5 (threshold for spike detection), cut_off = 500 (cut-off frequency for the butterworth filter in Hz). Automated clustering was followed by manual inspection, merging of units if necessary and discarding of noise and multi-units using phy (https://phycontrib.readthedocs.io). Units were evaluated based on the average waveform shape and auto-correlogram. Only cells without any inter-spike intervals of ≤ 1 ms were considered.

#### Analysis of in-vivo recordings

Unless otherwise noted, firing rates were calculated as the number of spikes in 50 ms bins averaged across the 10 stimulus repetitions. Z-scores were calculated as the number of standard deviations from the mean spontaneous activity before stimulus onset. All sorted units were grouped into cells with a maximal response amplitude > 2 standard deviations above the mean spontaneous firing rate (‘potentially responding,) and cells without such a peak (‘non-responding’). The activity to each stimulus repetitions was inspected for the ‘potentially responding’ cells to identify truly responding cells manually, which then were used for further analysis, average response calculations and visualization. For small stimuli shown at three different locations and moving in two different directions, only the strongest response was considered for population analysis.

##### DSI

Direction-selectivity was calculated based on the summed, back-ground subtracted activity during the time from the onset of the fast moving square until the end of the presentation for each direction α. These 8 response measurements R_k_ were normalized to the maximum and the DSI was calculated according to: 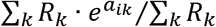.

##### Half-width of response to small, slow dot

Mean firing rates for each cell were background subtracted and the MATLAB function findpeaks was used to find the half-width of the highest peak.

#### Analysis of patch-clamp recordings

The loose-patch extracellular recording traces were high-pass filtered. Events that exceeded an amplitude threshold were extracted. Unless otherwise noted, firing rates were calculated as the number of spikes in 50 ms bins averaged across the 5-10 stimulus repetitions.

##### Chirp

Average responses were calculated based on the mean number of spikes during the stimulus across 10 trials.

##### Spot-size tuning curve

Firing rates were background subtracted and peak responses during the first 0.4 s after each stimulus onset were calculated and used to plot a spot-size tuning curve.

##### DSI

Direction selectivity was calculated as for the *in-vivo* recordings. Firing rates were background subtracted and peak responses during the first 1 s after each stimulus onset were calculated. The direction selectivity of a ganglion cell was defined as the vector sum of these peak responses for each of the 8 different directions α. These 8 response measurements R_k_ were normalized to the maximum and the DSI was calculated according to: 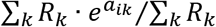.

#### Comparison of in-vitro and in-vivo data

To compare the response properties of different retinal ganglion cell types and neurons in the Pbg and pulvinar, we calculated z-scores for each responding neuron as described above. Median firing rates were plotted for the different brain nuclei and retinal ganglion cell types.

##### Latency measurements

For cells responding during the black expanding dot, firing rates were smoothed with a moving median and a window size of 150 ms. Then, the time to the maximal response amplitude was taken as a measurement for latency.

#### Cell body size measurements

To separate the functional group *xi* (ON alpha sustained cells) from group *x* (ON sustained and ON mini), we acquired a z-stack of most patched cells using the 2-photon setup. We then loaded the z-stack into Fiji, calculated a maximal projection and used the ellipse tool to fit an ellipse to the cell body and measure its area.

#### Statistics

To compare dendritic tree diameter distributions, we applied the Kolmogorov-Smirnov test (kstest2 function in MATLAB). Medians were compared by the Wilcoxon Ranksum test (ranksum function in MATLAB). We used both Pearson correlation and Spearman correlation (corr function in MATLAB) to test for significant gradients in the retinotopic distribution of dendritic tree diameters.

## Supplementary Information

**Figure S1 related to Figure 1:**
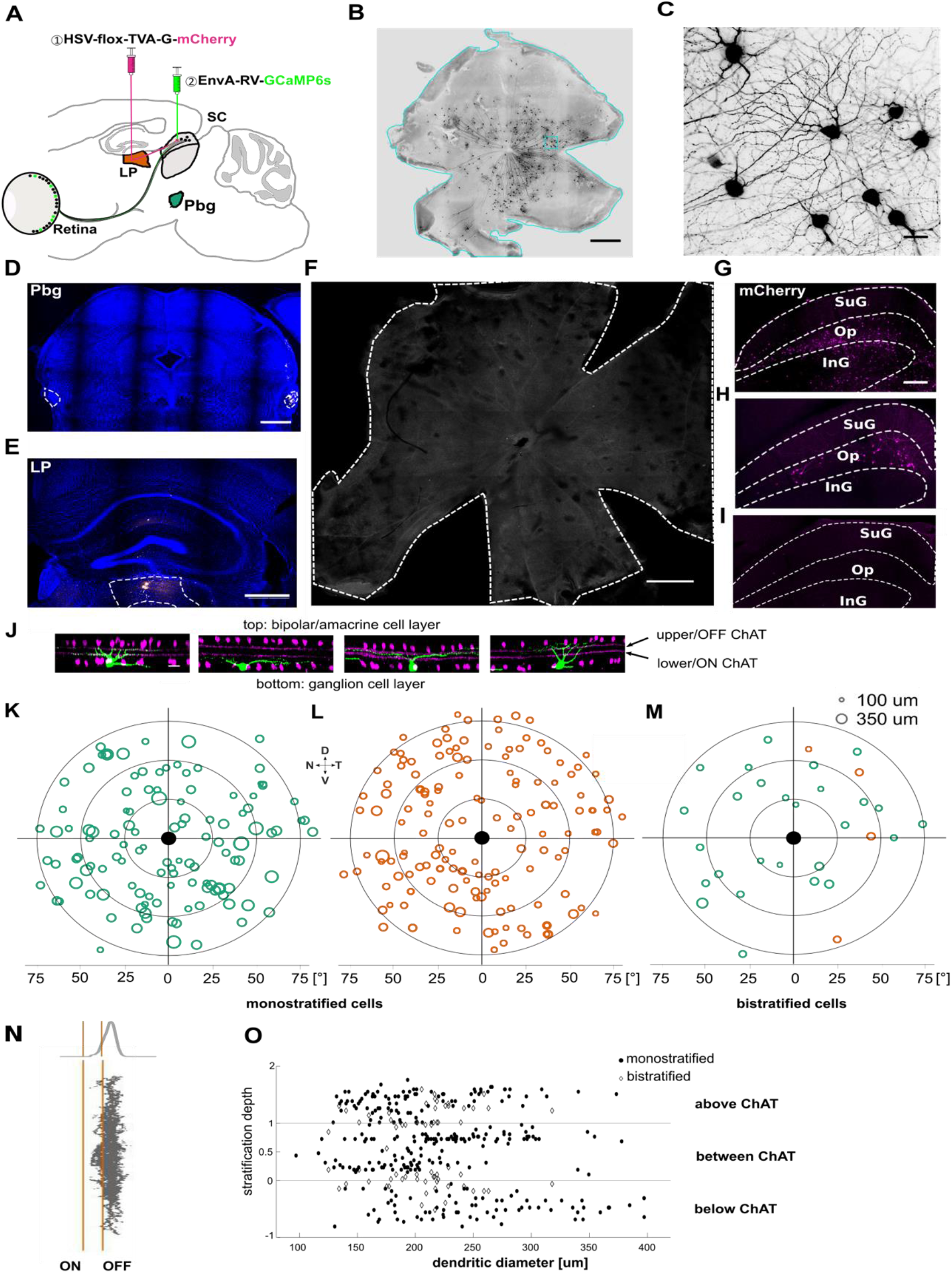
Viral tracing with EnvA-coated rabies virus and herpes-simplex-virus (HSV). **A)** Injection strategy for labelling of the circuit connecting the retina to the parabigeminal nucleus, via the superior colliculus. **B)** Example retina with labelled ganglion cells innervating the colliculo-parabigeminal circuit. Scale bar = 500 μm. **C)** Zoomed-in version of C. Scale bar = 50 μm. **D)** Histological section after parabigeminal nucleus injection. The pipette was coated with a fluorescent dye (DiD), and the fluorescent signal coincides with the location of the parabigeminal nucleus indicated with a dashed box. **E)** Histological section after pulvinar injection. **F)** The whole-mount retina stained with antibody for GCaMP6s after EnvA-coated SAD-ΔG-GCaMP6s rabies virus injection to superior colliculus alone without first injection of HSV. No labelled cells are observed after 11 days injection. Scale bar: 500μm. **G)** Injection of non-conditional HSV to parabigeminal nuclei labelled superior colliculus neurons. Neurons were stained with anti-mCherry antibody, showed in magenta. **H)** Injection of conditional HSV to pulvinar labelled superior colliculus neurons. Neurons were stained with anti-mCherry antibody, showed in magenta. **I)** Injection of conditional HSV to wild-type mouse. Very few labelled cells are observed after 21 days injection. Scale bar = 200μm. **J)** Side-view of z-stack scans of four example retinal ganglion cells (green) and the ChAT-bands (magenta). Scale bar = 20 μm. **K-M)** Retinal position and dendritic tree diameter of retinal ganglion cells that are part of the colliculo-parabigeminal circuit (K), cells innervating the colliculo-pulvinar circuit (L), and bistratified ganglion cells of both circuits (M). To determine if the differences in size between the colliculo-Pbg and colliculo-pulvinar circuit are due to a bias in the retinotopic location of the sampled ganglion cells, we analysed the spatial distribution of the labelled neurons across the retina. For each circuit we sampled evenly from each retinal quadrant (21.9% naso-dorsal, 26.6% dorso-temporal, 24.9% temporo-ventral, 26.9% ventro-nasal. In addition, we sampled at all retinal eccentricities: 16% of labelled ganglion cells were sampled from the central third of the retina (within 30° of the optic nerve), 48% from the middle third (30°-60° from the optic nerve) and 36% from the peripheral third (60°-90° from the optic nerve). This indicates that the observed difference in size between the two circuits is not due to a sampling bias in retinotopic location. N = nasal, D = dorsal, T = temporal, V = ventral. The optic nerve is indicated with a black disc. **N)** The distribution of the dendritic tree in depth was summed to create a stratification profile. **O)** Stratification depth and dendritic tree diameter of all 301 labelled retinal ganglion cells from both experimental conditions.

**Figure S2 related to Figure 2.**
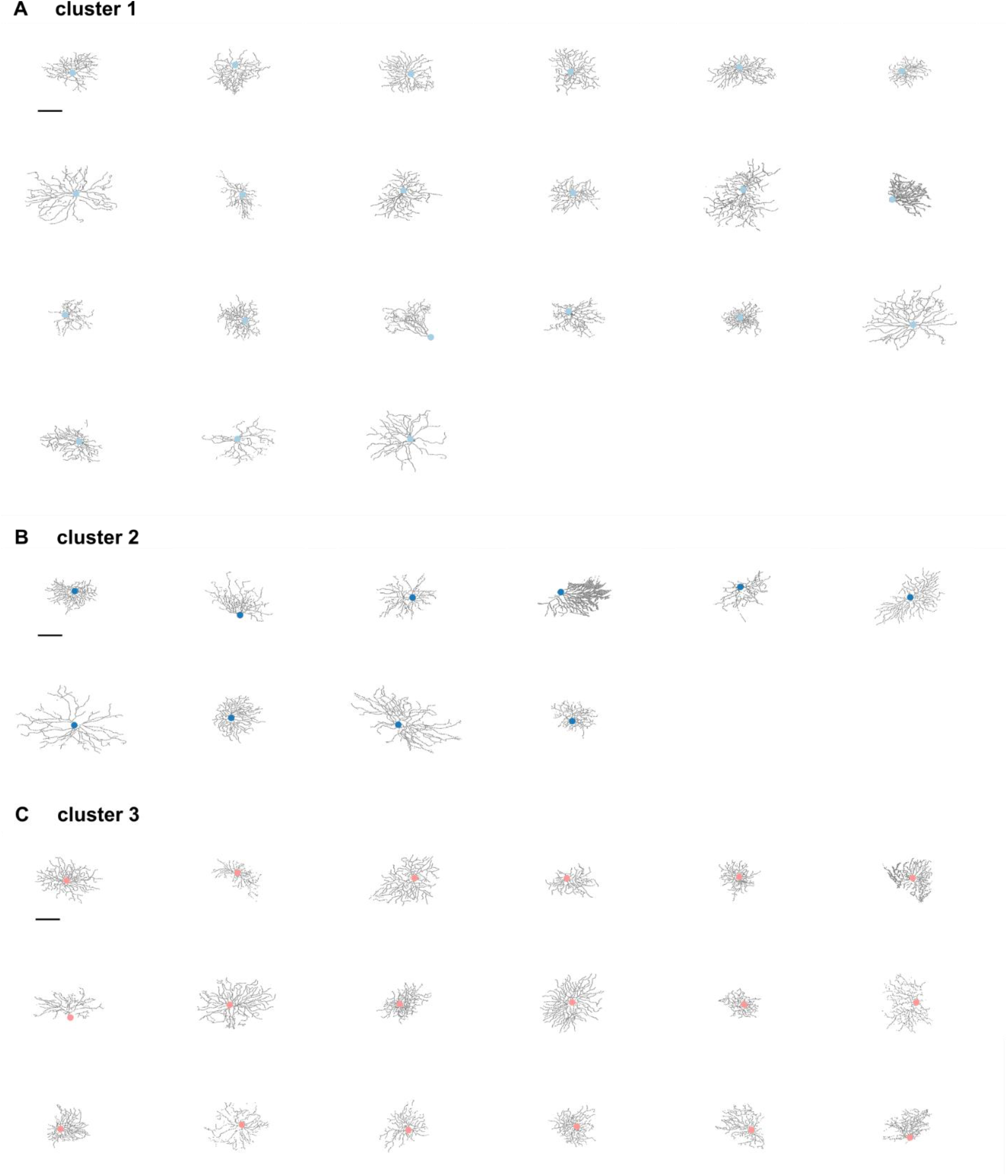
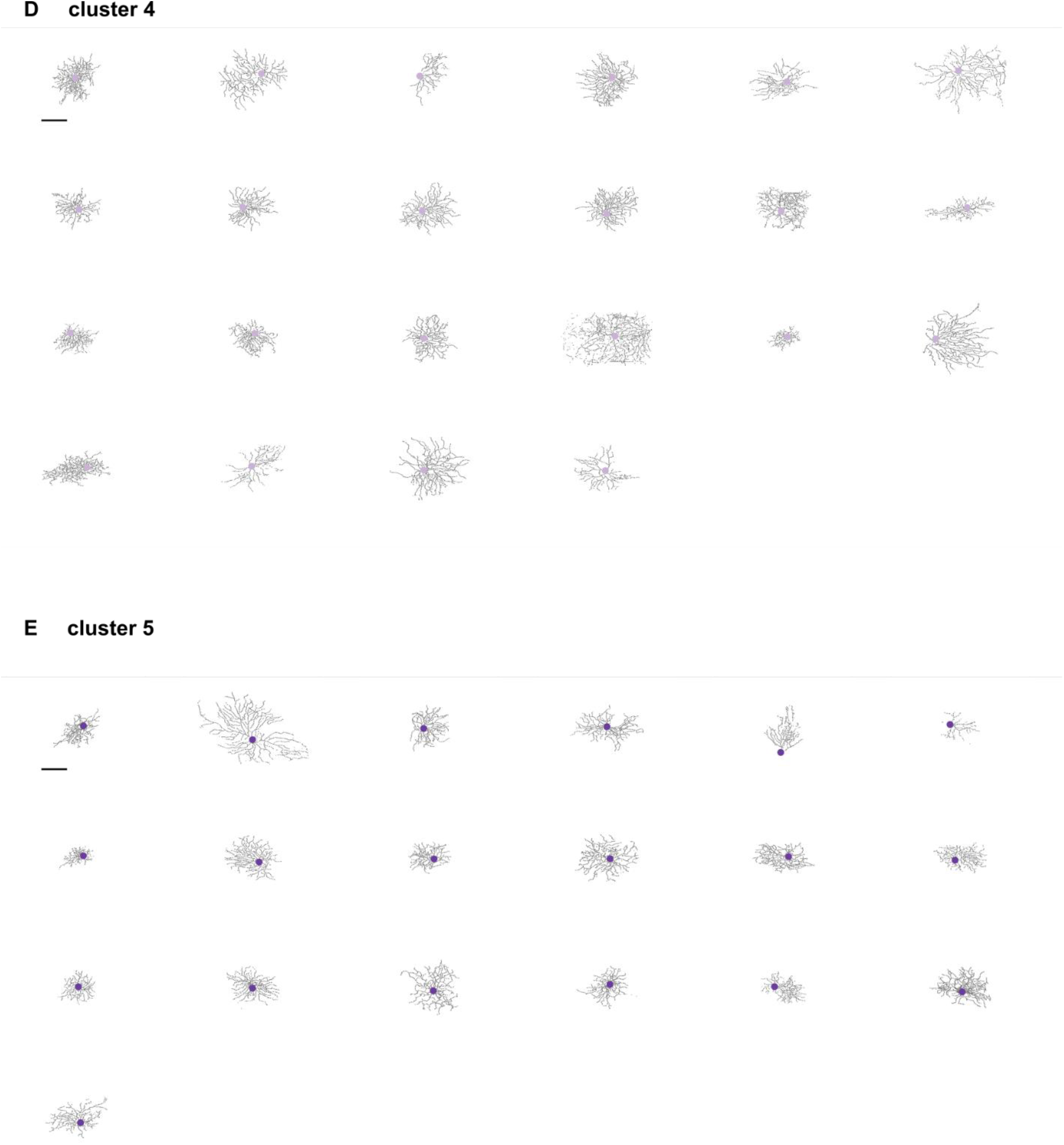
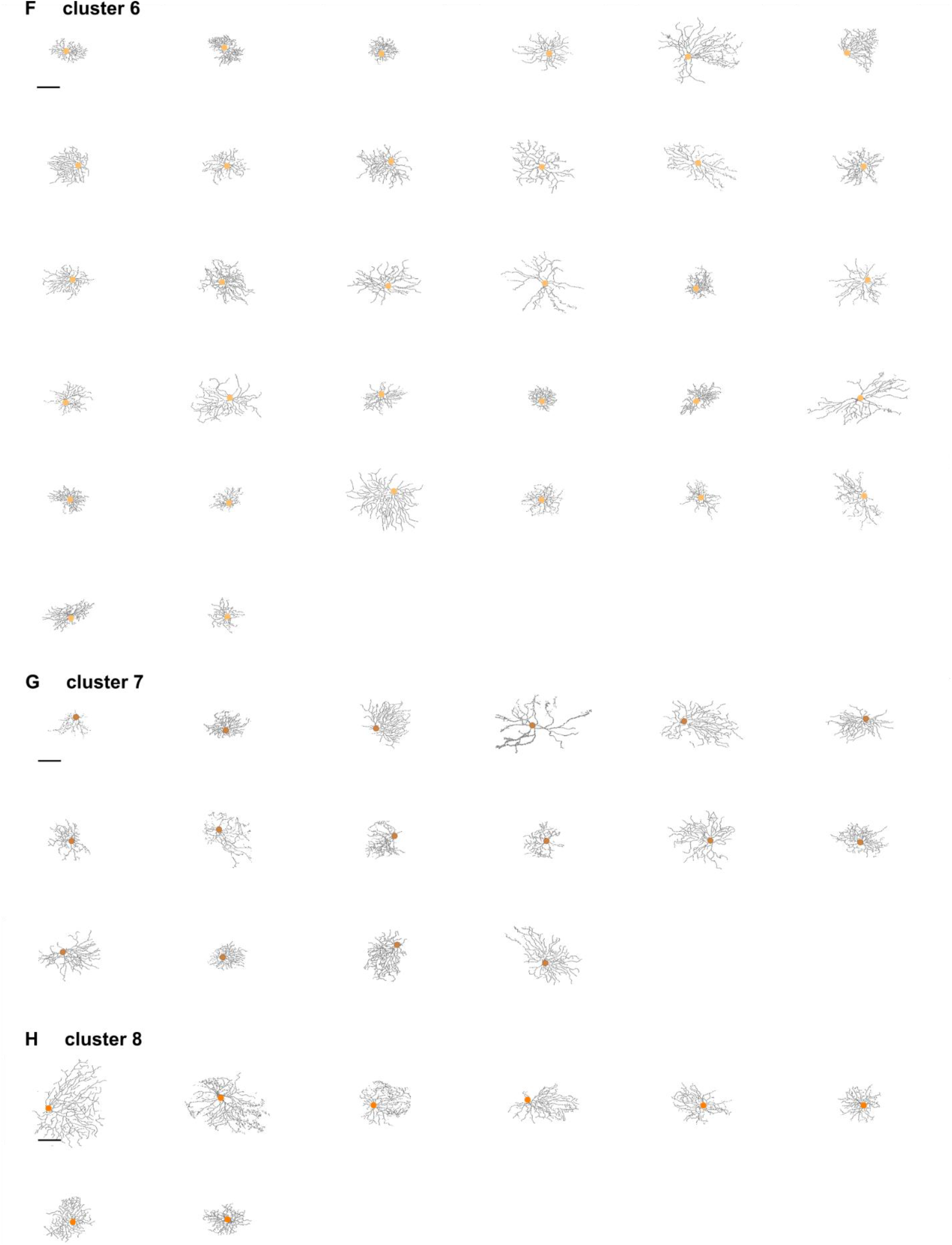
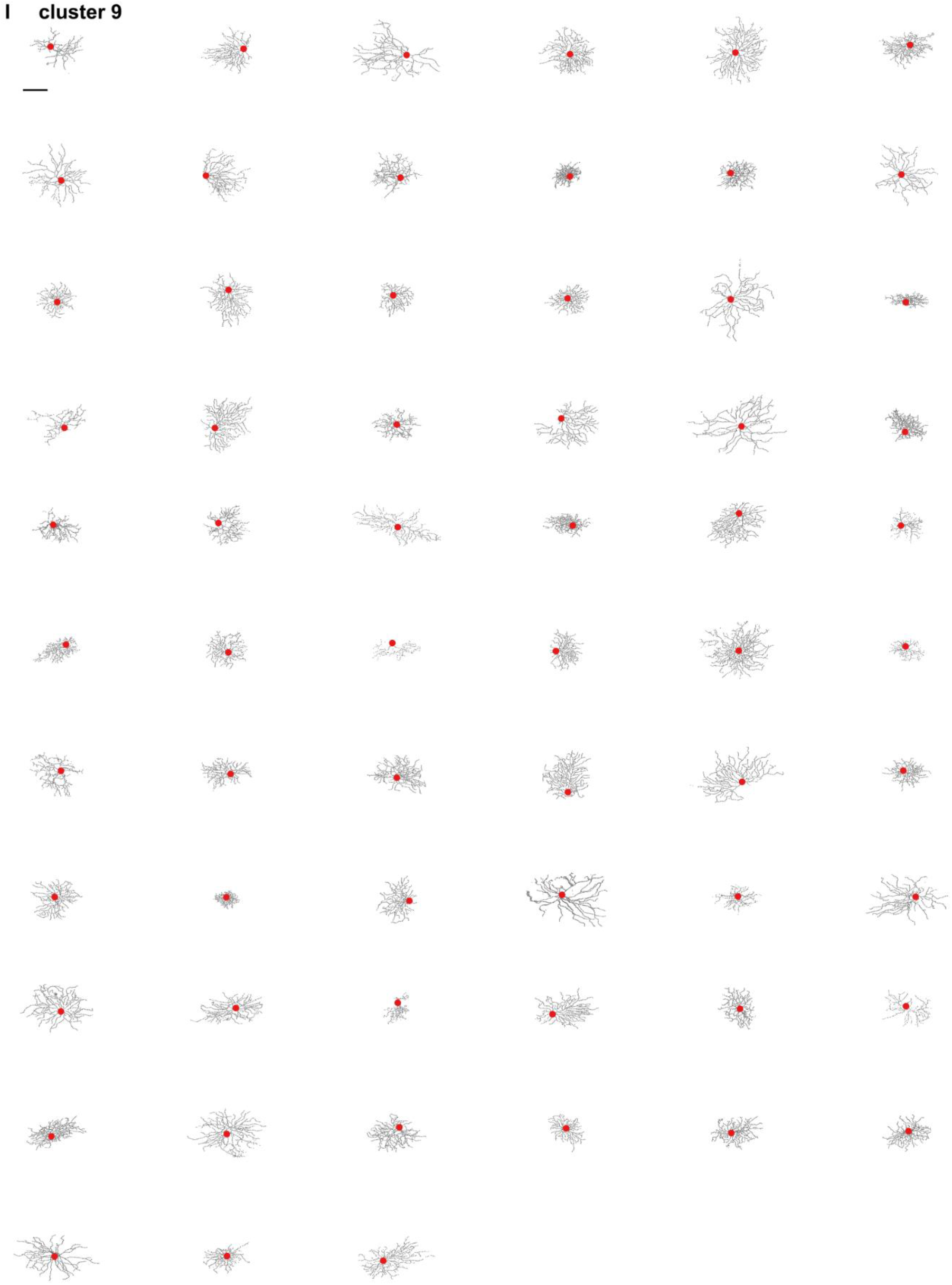
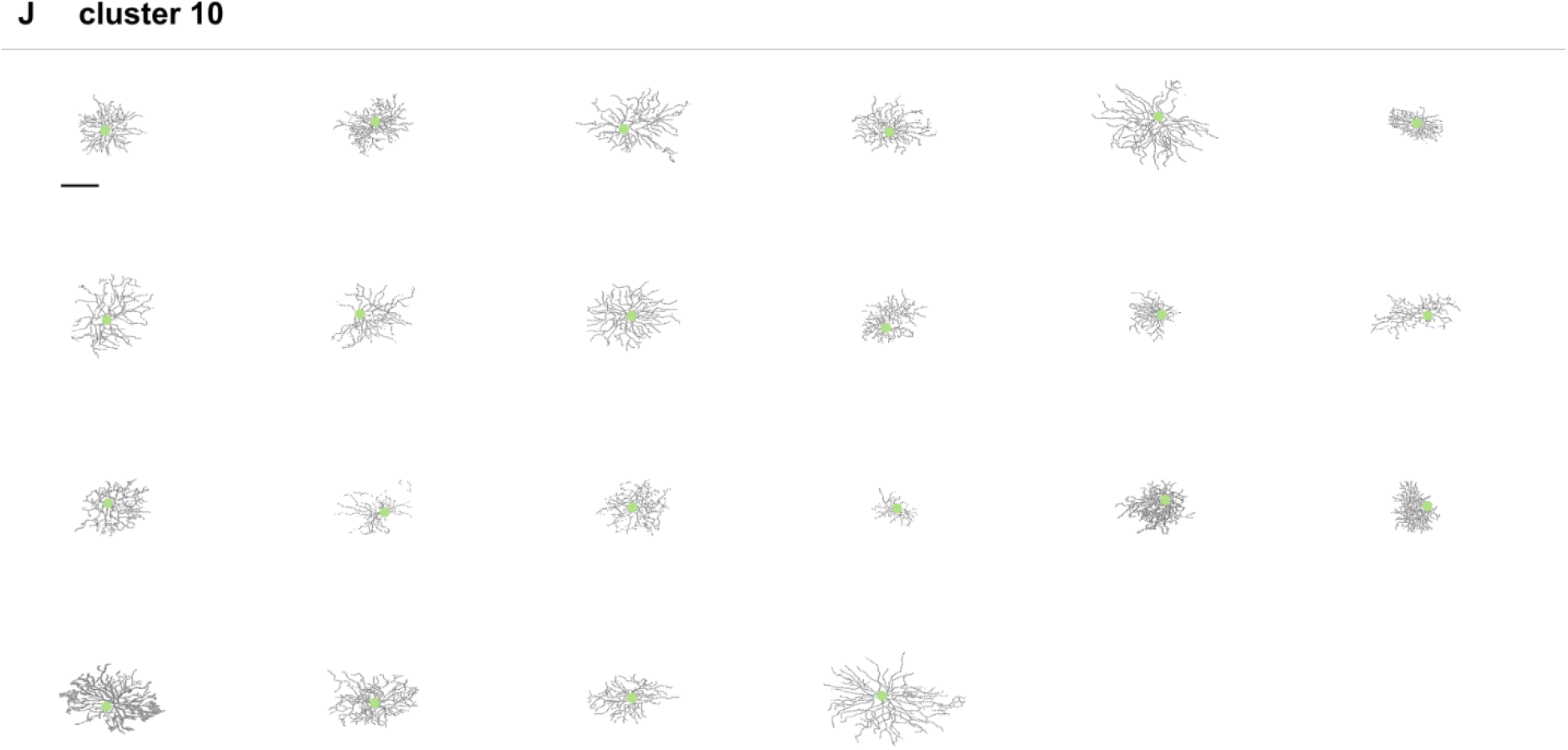
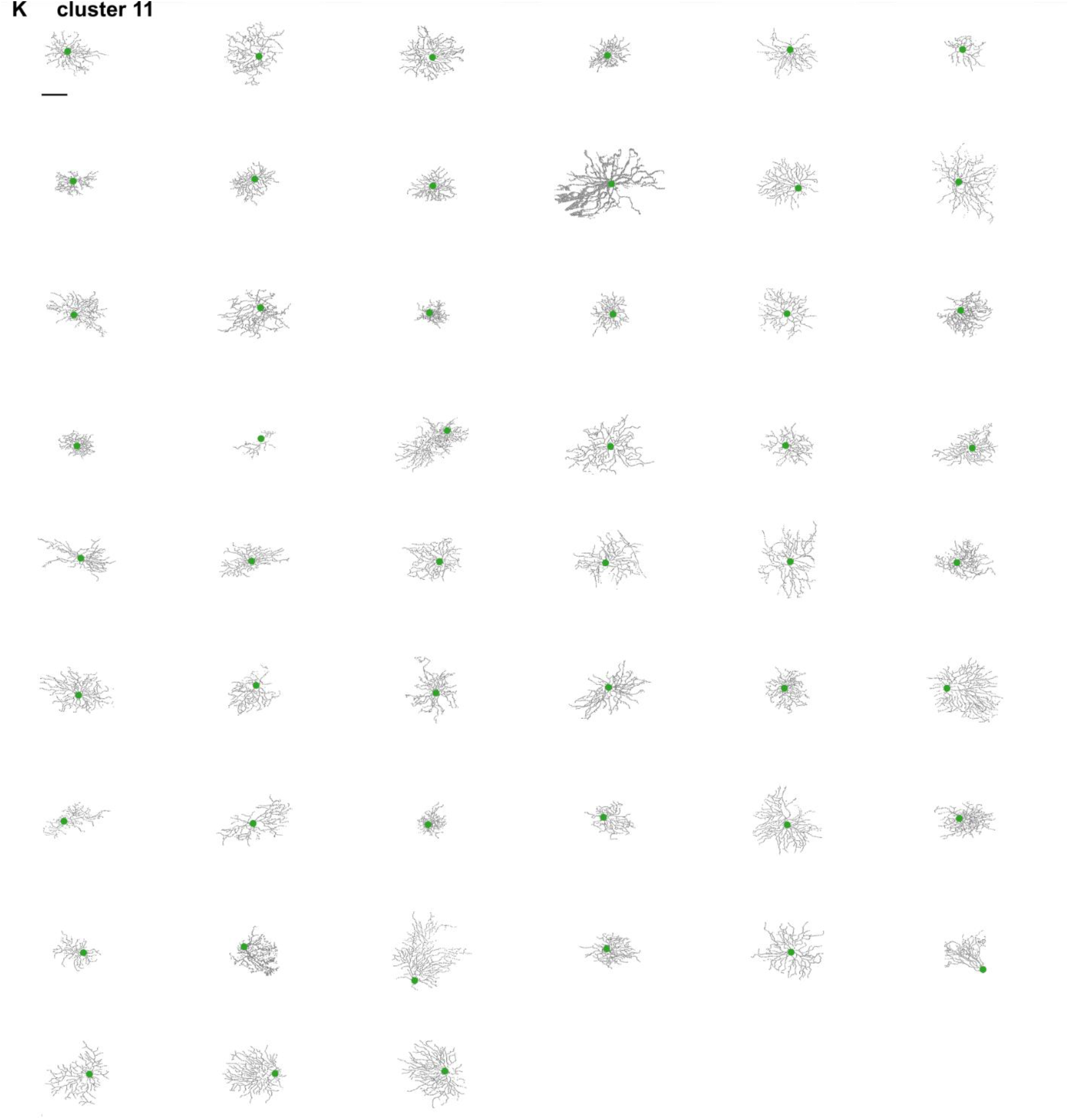
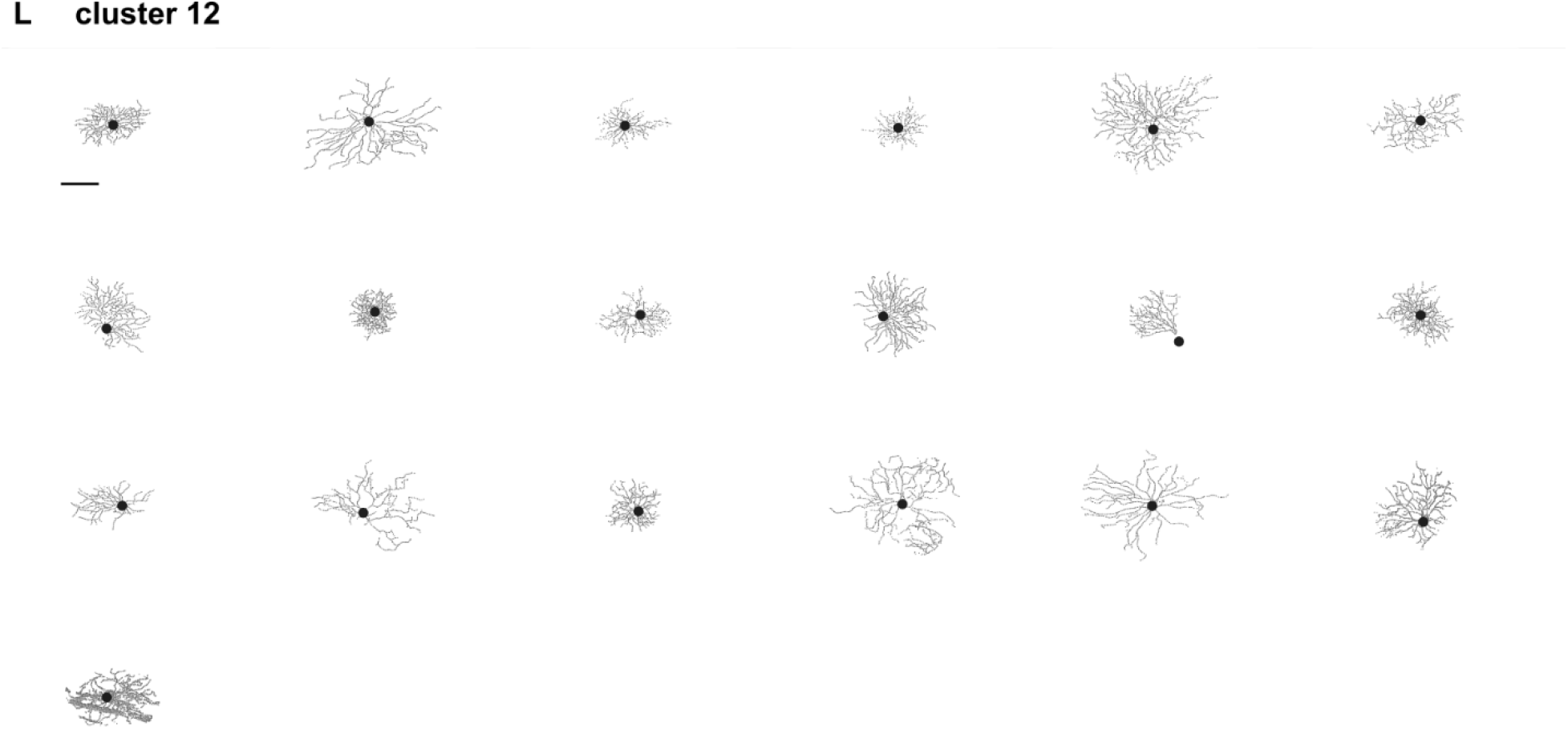
All 301 cells in their corresponding cluster. Cells are sorted by their similarity to the cluster centre (from closest do most distant). Scale bar: 100 μm.

**Figure S3 related to Figure 3.**
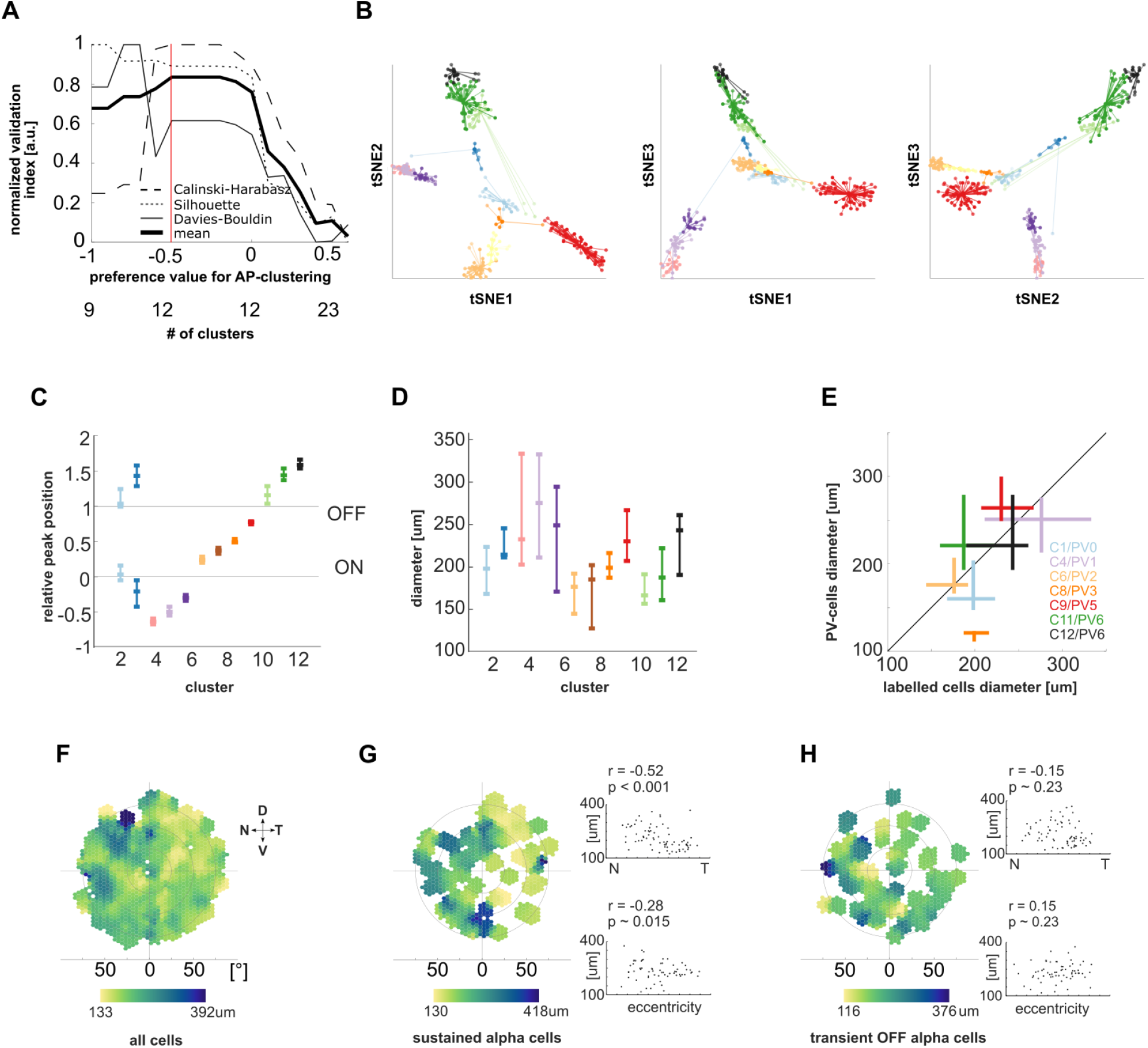
Clustering validation and dendritic tree size variability. **A)** Validation of clustering result. Three normalized validation indices (Calbinski-Harabasz, Silhouette, Davies-Bouldin) and their mean are shown. The clustering result taken for further analysis is marked with a red line. **B)** Visualization of the cells in the 12 clusters using t-distributed stochastic neighbor embedding (tSNE). Each plot shows the distances between individual data points. Transparency of the lines shows the similarity between two data points based on the affinity-propagation clustering. Colours represent the same clusters as shown in Figure 3. To test if an individual cluster contains retinal ganglion cells of more than one type, we compared the dendritic tree location and size variances within each group. **C)** Peak of stratification profile for each cluster (median and quartiles). **D)** Dendritic tree diameter for cells in each cluster (median and quartiles). The dendritic field size can vary substantially between cells of a given cluster, where generally smaller cells have a narrower range of dendritic diameters (e.g. cluster 6 and cluster 10), while larger cells span a broader range of sizes (e.g. cluster 5 and cluster 12). **E)** Cell diameter median and quartiles of 7 clusters and the corresponding parvalbumin-positive (PV) cells (Farrow et al., 2013) that have a similar stratification depth and similar average size. **F)** Smoothed distribution of dendritic field diameter of all labelled cells (n = 301) at their retinotopic location. To further investigate whether the within cluster variance in dendritic field size match the expected retinotopic distribution, we compared the size distribution of alpha-cell clusters with those previously reported (Bleckert et al., 2014; Krieger et al., 2017). When considering all 301 labelled cells, we find that cells in the central retina tend to be smaller than in the periphery. **G)** Smoothed distribution of dendritic field diameter of all cells in the sustained alpha clusters (n = 22 sustained ON-cells in cluster 4 and n = 51 OFF-cells in cluster 11), similar to published distributions (Bleckert et al., 2014). Top right: dendritic field size along the naso-temporal axis. Spearman-correlation: r = -0.52, p < 0.001; Pearson-correlation: r = - 0.42, p ~0.002. Bottom right: dendritic field size relative to eccentricity (from optic nerve to periphery). Spearman-correlation: r = 0.28, p ~ 0.015; Pearson-correlation: r = -0.19, p ~ 0.11. **H)** Smoothed distribution of dendritic field diameter of the cells in the transient OFF-alpha cluster (n = 63 in cluster 9). The size distribution of transient OFF-alpha cells is much more homogeneous and centred than for sustained alpha cells. Top right (naso-temporal axis): Spearman-correlation: r = -0.15, p ~0.23; Pearson-correlation: r = -0.14, p ~ 0.27. Bottom right (eccentricity): Spearman-correlation: r = 0.15, p ~0.23; Pearson-correlation: r = 0.14, p ~ 0.27.

**Figure S4 related to Figure 5:**
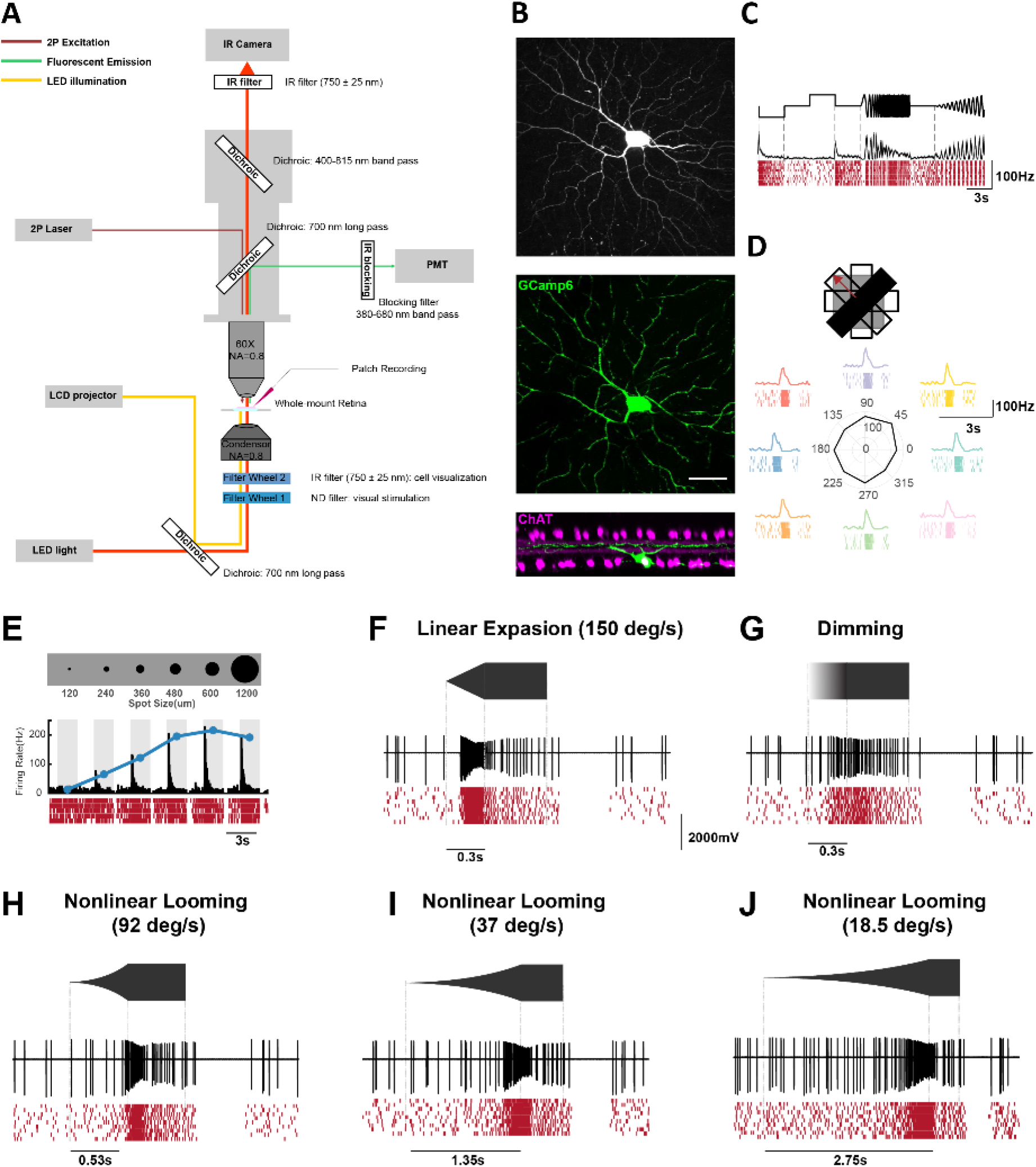
Targeted patch-clamp recording of the virus-labelled retinal ganglion cell. **A)** Schematic of the setup for the two-photon targeted patch-clamp recording. **B)** Top: Maximum intensity projection of a two-photon image stack showing GCaMP6-expressiong cell after the rabies injection. En-face view (middle) and side-view (bottom) of a confocal microscope z-stack (maximum intensity projection) showing the same cell after the staining process (green: GCaMP6, magenta: ChAT). Scale bar: 50 μm. **C)** Response of this cell to the chirp stimulus. The black trace representing the mean firing rates (50 ms bins) across 10 trials, which are shown below in the raster plot (red). **D)** Response to the black fast moving bar. **E)** Response to spot stimuli consisting of a black spot presented for 2s with 120, 240, 360, 480, 600 and 1200 μm diameter. The grey bars indicate stimulus duration. The blue line represents the spot size tuning curve. **F)** Response to a linearly expanding black dot. **G)** Response to a large dot that linearly dimmed from grey to black. **H-J)** Responses to non-linearly expanding black dots at different speeds.

**Figure S5 related to Figure 6:**
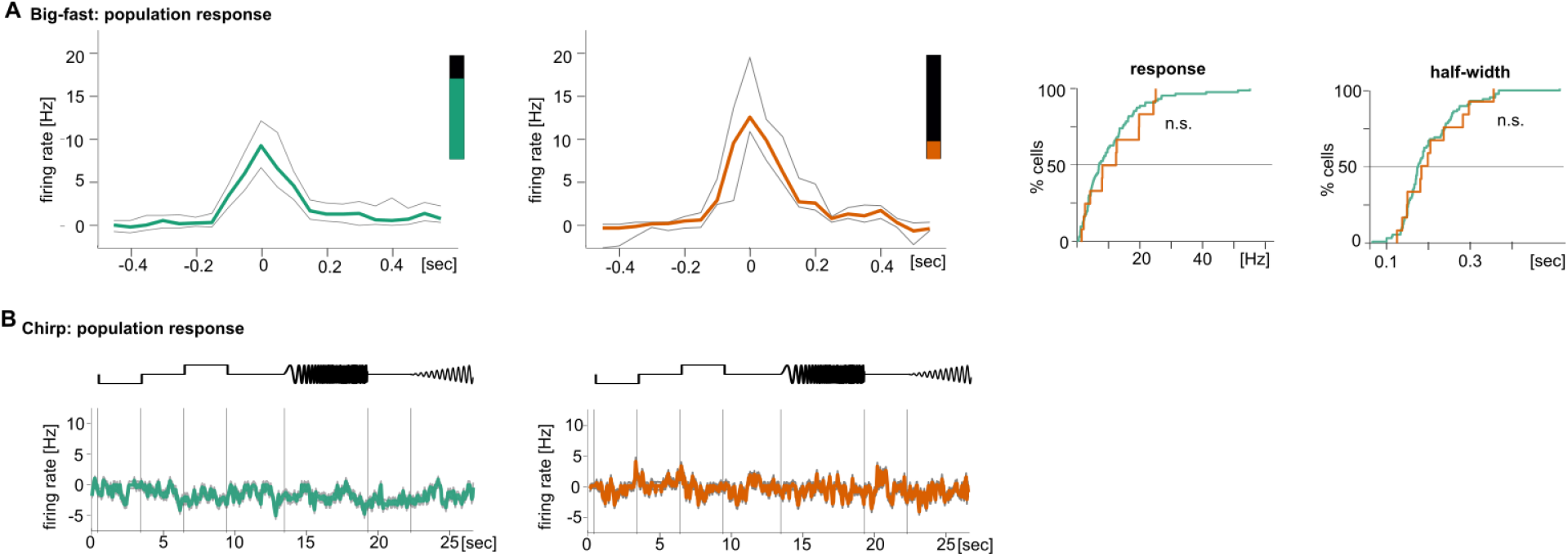
Parabigeminal and pulvinar responses to big-fast and chirp stimuli. **A)** Maximal amplitude response across all Pbg and pulvinar neurons to a big, fast moving black square. Right: Response amplitude distribution and half-width distribution. **B)** Population response to chirp stimulus.

**Figure S6 relates to Figure 7:**
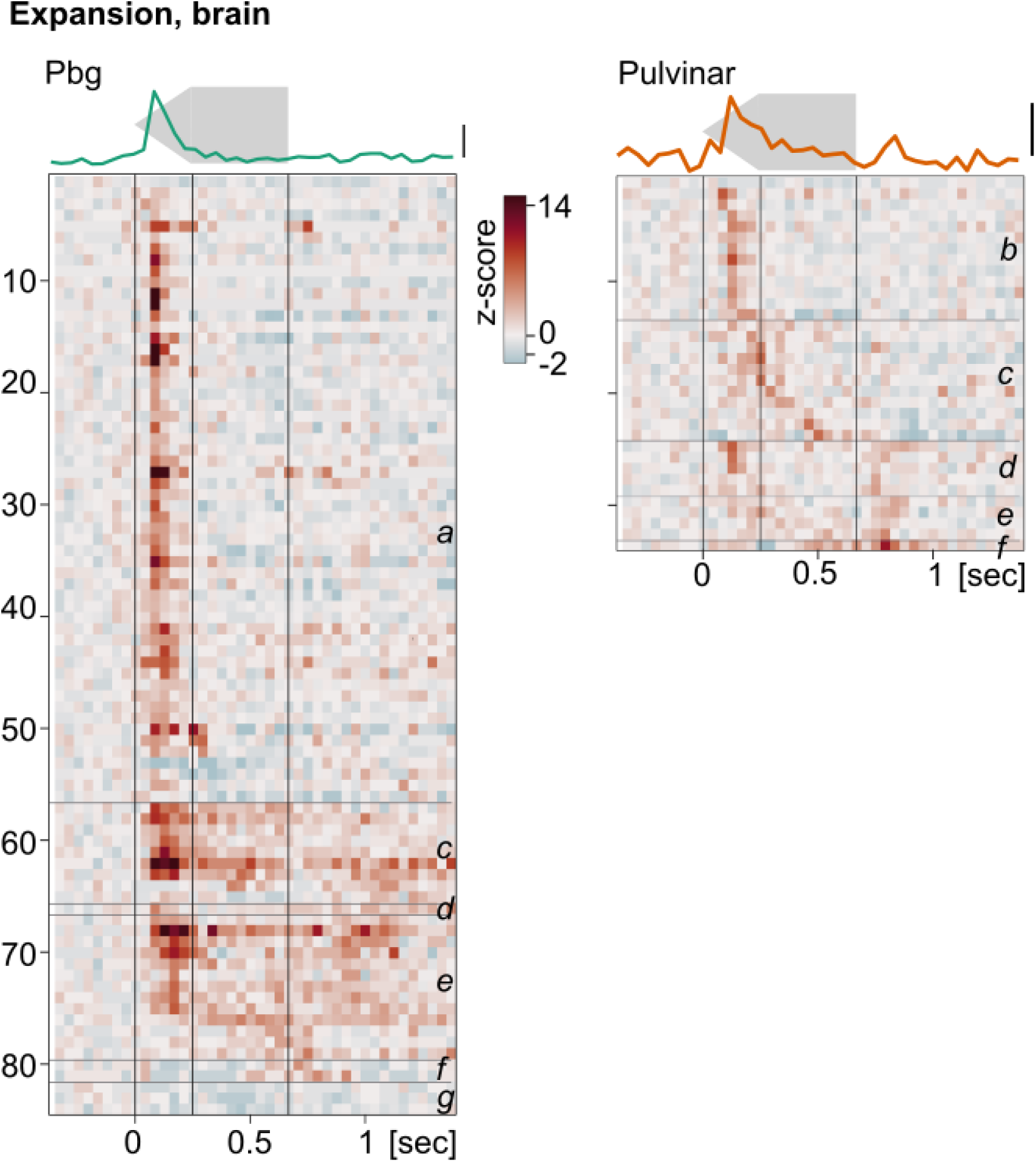
Parabigeminal and pulvinar responses to expansion. Heatmap of single cell responses to an expanding dot for parabigeminal nucleus and pulvinar. Letters indicate cells responding to a) onset; b) expansion; c) full size; d) expansion and offset; e) expansion, full size, and offset (sustained); f) offset only; g) expansion but with inhibition. Top: Average population response. Scale bar indicates z-score of 2.

**Table S1 related to Figure 7:**
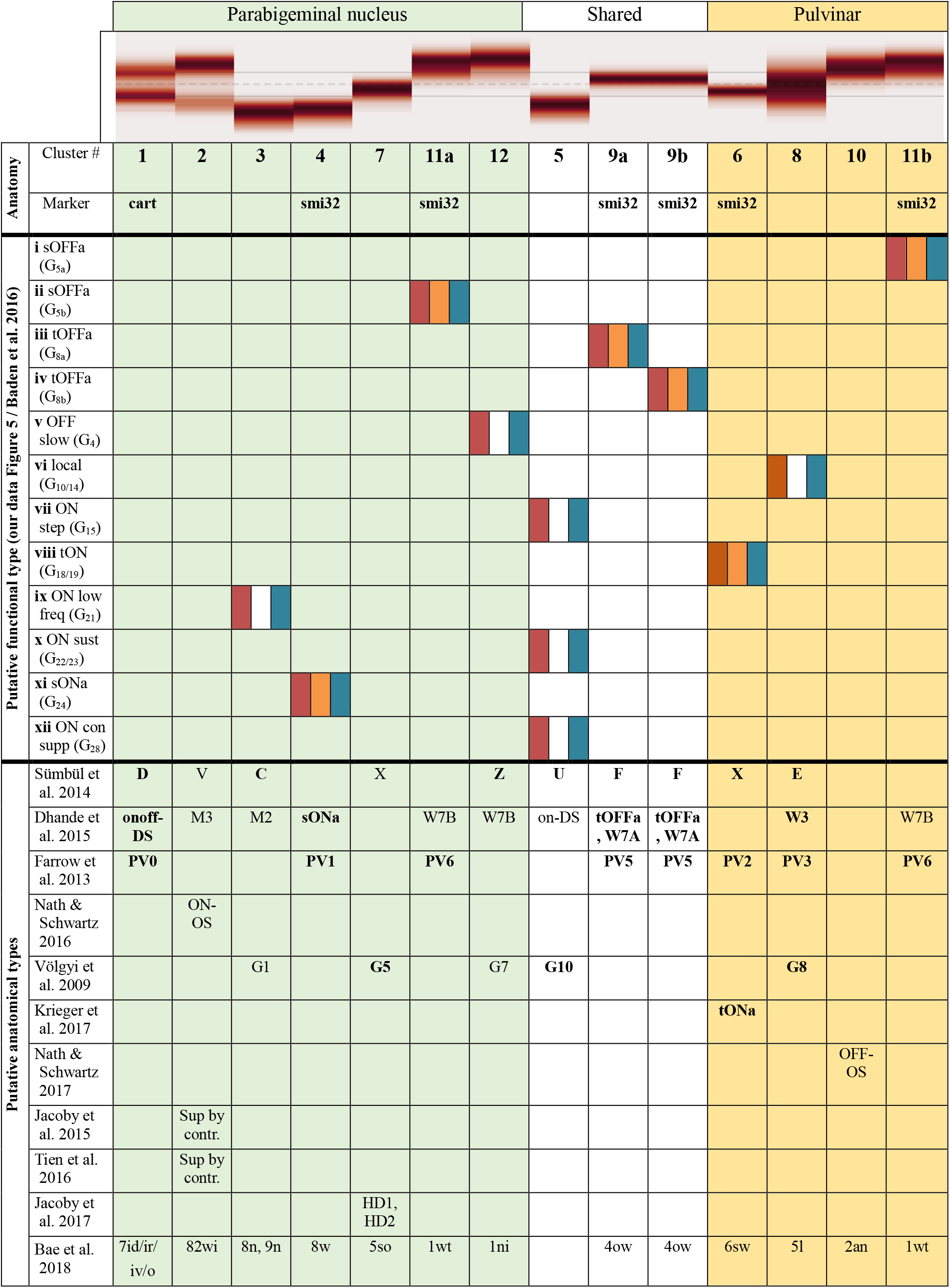
Putative retinal ganglion cell types of the colliculo-pulvinar and colliculo-parabigeminal circuit. Columns consist of anatomically identified cell types. Molecular markers, putative corresponding physiological clusters from our recordings, and putative anatomical cell types from the literature are given for each cluster. The pairing of anatomical and functional clusters is based on anatomy (red), molecular identity (orange), functional properties (blue) or a combination of those. The most likely putative anatomical types are highlighted in bold. ^1^Sümbül et al. 2014; ^2^Dhande et al. 2015; ^3^Farrow et al. 2013; ^4^Baden et al. 2016; ^5^Nath & Schwartz et al. 2016; ^6^Vöglyi et al. 2009; ^7^Krieger et al. 2017; ^8^Nath & Schwartz 2017

